# Lasting dynamic effects of the psychedelic 2,5-dimethoxy-4-iodoamphetamine ((±)-DOI) on cognitive flexibility

**DOI:** 10.1101/2023.07.05.547771

**Authors:** Merima Šabanović, Alberto Lazari, Marta Blanco-Pozo, Jason P. Lerch, Mark E. Walton, David M. Bannerman

## Abstract

Psychedelic drugs can aid fast and lasting remission from various neuropsychiatric disorders, though the underlying mechanisms remain unclear. Preclinical studies suggest serotonergic psychedelics enhance neuronal plasticity, but whether neuroplastic changes can also be seen at cognitive and behavioural levels is unexplored. Here we show that a single dose of the psychedelic 2,5-dimethoxy-4-iodoamphetamine ((±)-DOI) affects structural brain plasticity and cognitive flexibility in young adult mice beyond the acute drug experience. Using *ex vivo* magnetic resonance imaging, we show increased volumes of several sensory and association areas one day after systemic administration of 2mgkg^−1^ (±)-DOI. We then demonstrate lasting effects of (±)-DOI on cognitive flexibility in a two-step probabilistic reversal learning task where 2mgkg^−1^ (±)-DOI improved the rate of adaptation to a novel reversal in task structure occurring one-week post-treatment. Strikingly, (±)-DOI-treated mice started learning from reward omissions, a unique strategy not typically seen in mice in this task, suggesting heightened sensitivity to previously overlooked cues. Crucially, further experiments revealed that (±)-DOI’s effects on cognitive flexibility were contingent on the timing between drug treatment and the novel reversal, as well as on the nature of the intervening experience. (±)-DOI’s facilitation of both cognitive adaptation and novel thinking strategies may contribute to the clinical benefits of psychedelic-assisted therapy, particularly in cases of perseverative behaviours and a resistance to change seen in depression, anxiety, or addiction. Furthermore, our findings highlight the crucial role of time-dependent neuroplasticity and the influence of experiential factors in shaping the therapeutic potential of psychedelic interventions for impaired cognitive flexibility.

## Introduction

Serotonergic psychedelic drugs are psychoactive substances that induce acute changes in the perception of self, environment, and time, causing a non-ordinary state of consciousness, while also promoting long-term improvements in mood and psychological wellbeing [1, 2]. Growing evidence supports the lasting therapeutic effects of psychedelic-assisted psychotherapy for various conditions, including depression, anxiety, addictions, and personality disorders [3–10]. Healthy individuals also report higher positive affect and better wellbeing, effects that can persist for months [11] or up to a year following a single [12] or multiple treatment course [13], but the underlying mechanisms of these rapid and long-lasting behavioural changes remain unclear.

Psychedelics are capable of rapidly promoting structural and functional neuroplasticity shortly after administration [14]. Significant epigenetic and gene expression changes occur within hours [15–17], followed by a striking increase in dendritogenesis and spinogenesis that peak within 72h and decline over a week [17–21]. Although these immediate effects diminish, some structural and functional adaptations that have occurred in this period can potentially persist for weeks after [18]. The emerging dominant theory suggests that these psychedelic-induced neuroplastic adaptations can lead to long-lasting changes in mood and learning [2]. The psychedelic-induced window of plasticity may not be the sole catalyst for behavioural shifts, but instead could be acting as a gateway that improves learning or adaptability in conjunction with the heightened influence of environmental factors [2, 22, 23]. However, despite extensive and well-replicated research on the neuronal plasticity effects of psychedelics [24–30], the long-term impact of psychedelics on cognitive and behavioural plasticity, and their interaction with environmental factors, remains unclear.

Putative effects of psychedelics on cognition are particularly relevant considering that the psychedelics’ therapeutic effects have been shown across various mental disorders characterized by rigid cognitive and behavioural patterns and compulsive traits. Examples include rumination and negative cognition in anxiety and depression [31], difficulties in attentional switching in post-traumatic stress disorder [32], and compulsive rituals in eating disorders, addiction, and obsessive-compulsive disorder [33, 34]. In all these cases, impairments in cognitive flexibility are indicated by the persistence of a maladaptive response and a limited exploration of novel response strategies. Cognitive flexibility, which enables adaptation of beliefs or thoughts when they are no longer optimal [35], has been associated with greater resilience to negative life events and stress [36], suggesting that enhancing cognitive flexibility may have a protective and remedial effect on maladaptive stress responses. Yet, despite the suggested benefits of psychedelics on cognitive flexibility [2, 37–39], such effects have not been thoroughly investigated.

Existing evidence has produced mixed results, with studies suggesting both improvements [40–45] and deficits [45, 46] in cognitive flexibility following psychedelic administration, while others have found no discernible effects [47]. However, most studies to date have primarily focused on simple compulsivity tasks and serial reversals, testing only in the acute or sub-acute drug phase, despite the evidence that response inhibition [48] and working memory [49–51] are diminished under the influence of psychedelics. This approach overlooks the enduring behavioural changes that characterize the unique therapeutic effects of psychedelics. Therefore, there is a pressing need to investigate cognitive changes that manifest in the days and weeks following psychedelic treatment to begin uncovering the lasting consequences of psychedelic intervention.

To understand how structural and functional changes in individual neurons could translate to changes in mood and behaviour, our focus was on exploring the critical knowledge gap of psychedelic-induced plasticity at higher levels of analysis. Our primary objectives were, first, to confirm the structural plasticity effects of psychedelics at the level of whole-brain regions, and, second, to test any putative improvement in cognitive flexibility. We combined a single dosing regimen of a psychedelic substituted amphetamine, 2,5-dimethoxy-4-iodoamphetamine ((±)-DOI), with a multi-step reversal learning task and brain imaging in the post-acute phase of drug action. We hypothesized that the previously reported neuronal plasticity effects of psychedelics would be robust enough to result in observable changes in regional brain volumes, as measured by magnetic resonance imaging (MRI) during the period of highest neuronal plasticity. Furthermore, we expected that the rich synaptic landscape created by psychedelic-induced plasticity would facilitate enhanced speed and/or accuracy of reversal learning following (±)-DOI treatment, as assessed by adaptation to rule changes in a decision-making task. By investigating enduring structural and cognitive effects of psychedelics, we aimed to shed light on the long-term consequences of psychedelic-induced neuronal plasticity, providing valuable insights into the potential therapeutic applications of these substances.

## Methods

See Supplementary Information for detailed methods.

### Animals

All experiments were conducted under the UK Animal (Scientific Procedures) Act 1986 and the Local Ethical Review Committee at the University of Oxford. Adult male C57BL/6J mice (Charles River, Kent, UK) were used for all experiments. Food and water were available *ad libitum* unless the animals were under water restriction for behavioural testing (Supplementary Information).

### Drug

(±)-DOI hydrochloride (Sigma-Aldrich, D101-10MG) was dissolved in 0.9% saline (Aqupharm® No. 1), the vehicle control in all experiments. Each animal received only one injection, administered intraperitoneally at 5mlkg^−1^. A 2mgkg^−1^ dose of (±)-DOI was used for all experiments as it resulted in near-peak acute psychedelic effects when administered in a novel testing environment (Fig.S1).

### Ex vivo MRI

The sample preparation procedure was adapted from previous reports (Supplementary Information) [52]. Images were acquired with a 7T field strength Bruker BioSpec® 70/20 USR multipurpose high field MR scanner with a receive-only CryoProbe (Bruker BioSpin) coil. T2-weighted turbo spin-echo rapid acquisition with relaxation enhancement (TSE RARE) sequence parameters were: 12ms echo spacing, 6 echoes, TR/TE=350/12ms, BW=60kHz, 400ξ160ξ200 matrix, 60µm isotropic voxels, imaging time 33min.

Image analysis and visualization were performed using the RMINC library in R [53]. An analysis of variance (ANOVA) for the main effect of the drug was computed across the whole brain, either at every voxel or every region of interest (ROI). To assess the relative differences across groups, *t*-tests were computed at every voxel/ROI comparing samples from (±)-DOI- and vehicle-treated mice. False discovery rate (FDR) was set at an explorative 20% to maximize discovery rate for unbiased whole-brain analyses.

### Two-step reversal learning task

Mice were trained on a two-step probabilistic reversal learning task, as described previously [54] (training timeline in Table S1). The task required water-restricted mice to initiate a trial by poking a central port and then choose between two possible step 1 ports (left/right) to access a water reward at one of the two step 2 ports (up/down). In step 2, one of the up/down reward ports lit up for the animal to poke and water reward was delivered according to a probabilistic schedule. Reward probabilities were anticorrelated – i.e., when reward probability was high (80%) for one port, it was low (20%) for the other – and reversed serially in each session. Each step 1 choice was associated with a *common transition* (80% of trials) to one step 2 reward port and a *rare transition* (20% of trials) to the other. Transitions were initially fixed for each mouse (counterbalanced across animals). During post-drug testing, a single transition reversal was implemented, either one week (*Two-step Experiments 1 & 3*) or one day (*Two-step Experiment 2*) after drug treatment (Fig.S2), making the previously common transitions rare, and vice versa.

### Data analysis

General task performance was quantified as the mean number of trials, correct choices, and reward reversals completed in one session. Trial-to-trial learning was assessed using a logistic regression model (Supplementary Information) predicting the likelihood of repeating a choice based on the earlier trial events (type of transition and outcome). We also implemented a second logistic regression that evaluated the effect of transition type on rewards and omissions separately. Significance of individual predictors was determined by one sample *t*-tests (or Wilcoxon signed-rank) against zero. Pre-drug performances were compared via unpaired *t*-tests, or two-way repeated measures (RM) ANOVA for pre-versus post-drug comparisons, supplemented by Bayesian equivalents.

Adaptation of choice strategies to reward reversals was assessed by calculating average choice probability in the first 20 post-reversal free-choice trials, averaging across individual subjects, and comparing across groups with a permutation test (Supplementary Information).

Adaptation of choice strategies to the transition reversal was assessed with the logistic regression models applied across three concatenated sessions to supply enough trials for accurate model fits. Time series of regression coefficients were then fitted using nonlinear regression. First, best-fit models (horizontal line, line, or one-phase association) were found for (±)-DOI and control datasets separately. The drug effect was considered significant if best-fit models differed across datasets, *or* if separate fits for the same best-fit model were preferred over a global fit across both datasets. The same approach was used to compare post-reversal general task performance over time. Detailed analysis methods are in Supplementary Information.

## Results

See figure legends for detailed statistical results.

### Grey matter volume changes one day after (±)-DOI

We first wanted to examine whether the previously reported (±)-DOI-induced increases in dendritic branching and spinogenesis [17, 19, 20] would be reflected in discernible changes in regional grey matter volume. We injected mice with either 2mgkg^−1^ (±)-DOI or saline vehicle and collected their brains 24-36h later (Fig.1A, *n*_group_=8). At an explorative 20% FDR threshold, hierarchical unbiased ROI-wise *ex vivo* MRI analysis returned significant enlargements of several sensory areas (Fig.1B-E, Fig.S3): primary and lateral secondary visual areas (V1 and V2L, respectively), ventral secondary auditory cortex (AuV), and parts of primary somatosensory cortex (S1). Additionally, transmodal association regions such as temporal association area (TeA), lateral parietal association area (LPtA), and retrosplenial agranular area (RSA) were significantly larger in (±)-DOI-treated samples. V1 (*q*=0.016) and TeA (*q*=0.043) survived the conservative 5% FDR threshold. Volume enlargements ranged from 16.4 ±3.9% (mean ±SEM) in the shoulder region of left S1 to 6.6 ±1.8% in the left RSA (Table S2). In each ROI, the left hemisphere was the one showing statistically significant differences, with the right hemisphere volumes not crossing the significance threshold.

**Fig.1.**
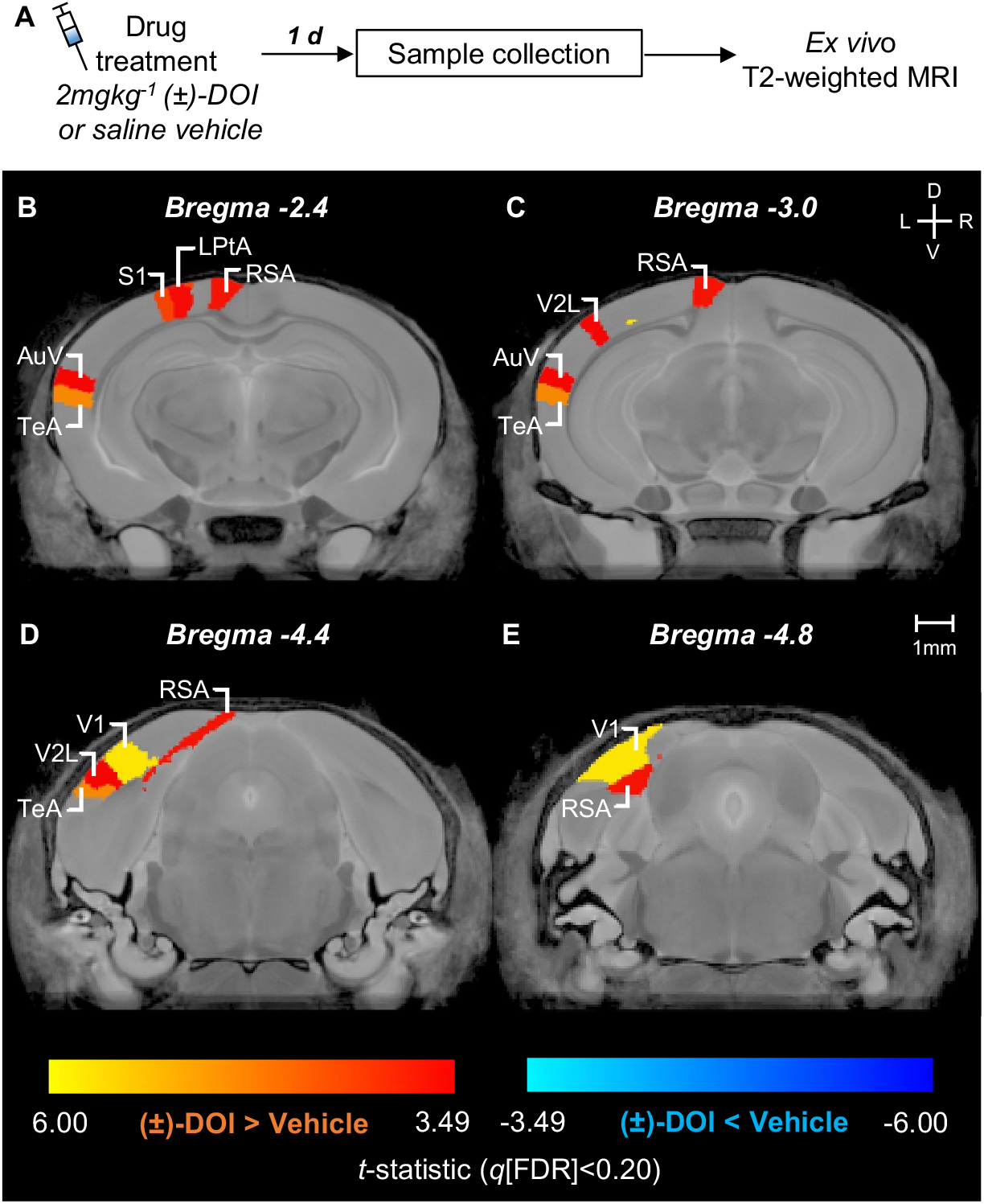
(±)-DOI increased the volume of several sensory and association cortical areas within one day. **(A)** Experiment timeline. The brains were collected one day after an injection of either 2mgkg^−1^ (±)-DOI or saline vehicle. **(B-E)** The False Discovery Rate (FDR) correction at explorative 20% resulted in the marginal *t*-statistic =3.49 (linear model *F*_1,14_=12.19). Significant increases were in primary somatosensory (S1) shoulder (*t*=4.25) and trunk (*t*=4.15) areas, retrosplenial agranular area (RSA, *t*=3.68), lateral parietal association area (LPtA, *t*=3.61), ventral secondary auditory area (AuV, *t*=3.53), and lateral secondary visual cortex (V2L, *t*=3.49). Volume increases in the primary visual area (V1, *t*=5.75) and temporal association area (TeA, *t*=4.83) survived the strict 5% FDR (*F*_1,14_=23.30, marginal *t*-statistic =4.83). *n*_group_=8. D: dorsal. V: ventral. L: left. R: right.

To further understand the spatial distribution of volume changes, we determined local volume differences with a more detailed, voxel-wise analysis. Significant enlargements at the 20% FDR threshold were also detected in parts of the right hemisphere V1, V2L, and S1 (Fig.S4). Significant voxel-wise differences were found in regions other than those highlighted in the ROI-wise analysis. Some of these were bilateral (e.g., lateral posterior thalamic and caudal pontine reticular nuclei), while others were restricted to the left (e.g., motor cortex and CA1) or right (e.g., anterior olfactory and amygdaloid areas, entorhinal cortex) hemisphere only. Taken together, these data demonstrate that (±)-DOI induced post-acute grey matter structural changes across cortical and subcortical areas.

### Pre-drug cognitive strategies in the two-step task

Given the suggested relationship between psychedelics and neuronal plasticity [17–21], we hypothesized that (±)-DOI could shape the synaptic landscape to be conducive to improved learning and cognitive flexibility [2, 38]. We explored the potential impact of (±)-DOI on cognitive flexibility by training young adult water-restricted mice (*N*=82) on a two-step reversal learning task in which subjects made a left/right choice in step 1 to access an up/down port in step 2 which delivers water reward on a probabilistic schedule (Fig.2A-B). To maximise reward rate, mice had to track the reward probabilities at up/down ports, which were anticorrelated and reversed in series, and the transition probabilities, determining the likelihood of an initial left/right choice leading to a particular up/down state, which were fixed (Fig.2B-C).

**Fig.2.**
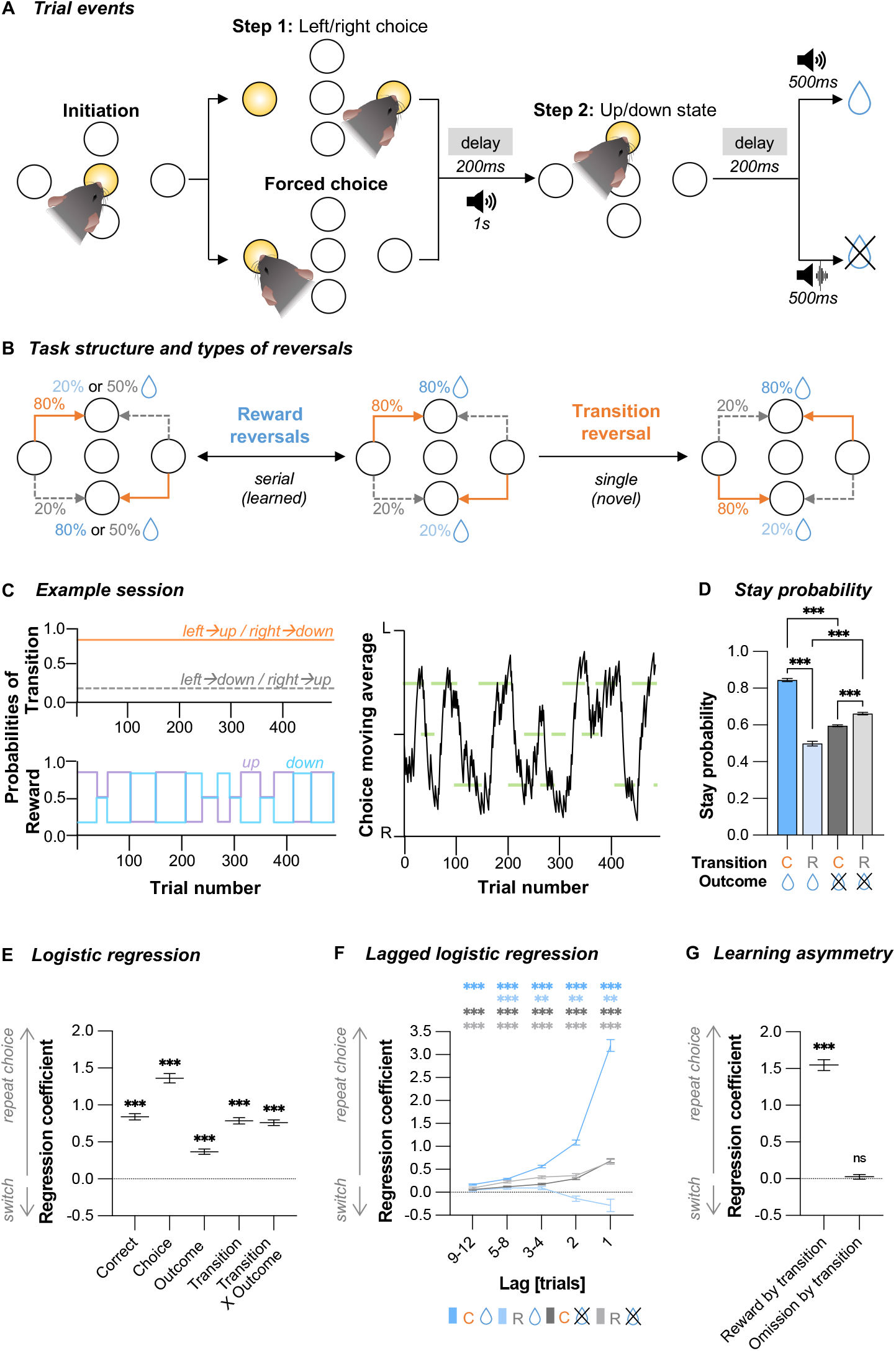
Choice performance on the two-step task prior to any drug treatment reflects the combined influence of outcomes and transition types. **(A)** Trial events. A mouse initiated the trial by poking the central port. In step 1, either both left/right ports lit up for the animals to choose which one they poke (*free choice*), or only the left or right port lit up to force the animal to explore that choice (*forced choice*, max. 25% of trials per session). Distinct auditory cues signalled active up or down port. A tone (identical to the up/down cues) or white noise would cue water reward delivery or omission, respectively. **(B)** The probabilistic structure of the task and the types of reversals. Reward probabilities of step 2 states reversed in blocks which could be non-neutral (reward probabilities switch between 80% and 20%) or neutral (both reward probabilities 50%). *Reward reversals* were triggered based on a behavioural criterion for non-neutral blocks (random interval of 5-15 trials after the exponential moving average across 8 previous free choices >75% correct) or after a random 20-30 trial interval for the neutral block. Animals were trained on serial reward reversals (“learned” adaptations). The transition structure was initially fixed until a single experimenter-directed *transition reversal* that occurred after drug treatment (“novel” adaptation). **(C)** Example sessions. Top: Transition and reward probabilities. Bottom: exponential moving average of choices (tau=8 trials). Green lines represent reward blocks, and their *y*-position represents the correct choice (left, right, or neutral). **(D)** Stay probabilities for the step 1 choice were a function of subsequent common (C) and rare (R) transitions and reward (+), or omission (−), trial outcomes, as well as their interaction. RM two-way ANOVA (all *P*<0.001, BF_incl_>>100): *Transition F*_1,81_=351.7, η_p_^2^=0.81; *Outcome F*_1,81_=43.9, η_p_^2^=0.35; *Transition* X *Outcome F*_1,81_=541.1, η_p_^2^=0.87. Stars represent Bonferroni-corrected post-hoc comparisons. **(E)** Logistic regression analysis quantifying how transitions, outcomes, and their interaction predict repeating the same step 1 choice on the next trial. *Correct* and *Choice* predictors correct for cross-trial correlations and capture any side bias. One sample *t*-test or Wilcoxon signed rank tests against zero (all *P*<0.001, BF_10_>>100): *Correct t*_81_=19.8, Cohen’s *d* =2.18; *Choice V*=3402, Rank-Biserial Corr. =1.00; *Outcome V* =3286, Rank-Biserial Corr. =0.93; *Transition V* =3403, Rank-Biserial Corr. =1.00; *Transition X Outcome V*=3403, Rank-Biserial Corr. =1.00. **(F)** Lagged regression analysis shows how the repeated step 1 choices were influenced by the trial history. Stars represent family-wise Bonferroni-corrected one sample *t*-tests or Wilcoxon signed rank tests against zero. **(G)** Mice exhibit asymmetry in how they learn from positive and negative feedback. The reinforcing value of rewards was modulated by the type of transition, but reward omissions did not contribute to trial-by-trial learning. One sample *t*-tests or Wilcoxon signed rank tests against zero: *Reward by transition V*=3403, *P*<0.001, Rank-Biserial Corr. =1.00, BF_10_>>100; *Omission by transition t*_81_ =0.83, *P*=0.411, BF_10_=0.17. Data shown as mean ±SEM. *n*=82. \**P*<0.05. \*\**P*<0.01. \*\*\**P*<0.001.

Pre-treatment, mice became proficient at the task in 27 ±4 (mean ±SD) training sessions, completing 388 ±71 trials and 10 ±3 reward reversals per session when fully trained. Analysing trial-by-trial choice strategies showed that mice repeated rewarded choices but, importantly, also exhibited sensitivity to the underlying transition structure (Fig.2D, Transition X Outcome *P*<0.001). The reinforcing effect of trial outcome depended on the common/rare transitions that preceded them (Fig.2E, *Transition X Outcome* predictor *P*<0.001). This effect persisted across multiple trials (Fig.2F). This pattern indicates mice understood the task structure and tracked which step 1 action commonly led to a particular step 2 state.

Consistent with previous work using the same task [54], transition type affected stay/switch behaviour following rewards but not omissions (Fig.2F). To verify this, we adapted the logistic regression analysis to evaluate the influence of transition type on rewarded and unrewarded trials separately (Fig.2G). As predicted, we found that common transitions on rewarded trials promoted staying (*Reward by transition* predictor *P*<0.001), but omission trials had no significant impact on stay/switch likelihood (*Omission by transition* predictor *P*=0.411), demonstrating asymmetry in learning from different trial outcomes.

### Unmasking the effects of (±)-DOI on cognitive flexibility

To evaluate post-acute effects of (±)-DOI on cognitive flexibility, in a first cohort (*Two-step Experiment 1*, *N*=26, Fig.S2), we injected fully trained mice with (±)-DOI or saline vehicle and, starting one day later, tracked their task performance over the course of one week (Fig.3A-B). Compared to the pre-drug period, there were marginal increases in the number of trials (Fig.S5A, Time *P*=0.018, mean difference ±SEM =16 ±6, Drug *P*=0.391) and reward reversals per session (Fig.3D, Time *P*=0.048, mean difference =0.7 ±0.3, Drug *P*=0.160) in both (±)-DOI and vehicle treatment groups, but there was no significant improvement in the number of correct choices (Fig.3C, Time *P*>0.521, Drug *P*=0.566), indicating that (±)-DOI did not enhance the animals’ overall decision-making accuracy.

**Fig.3.**
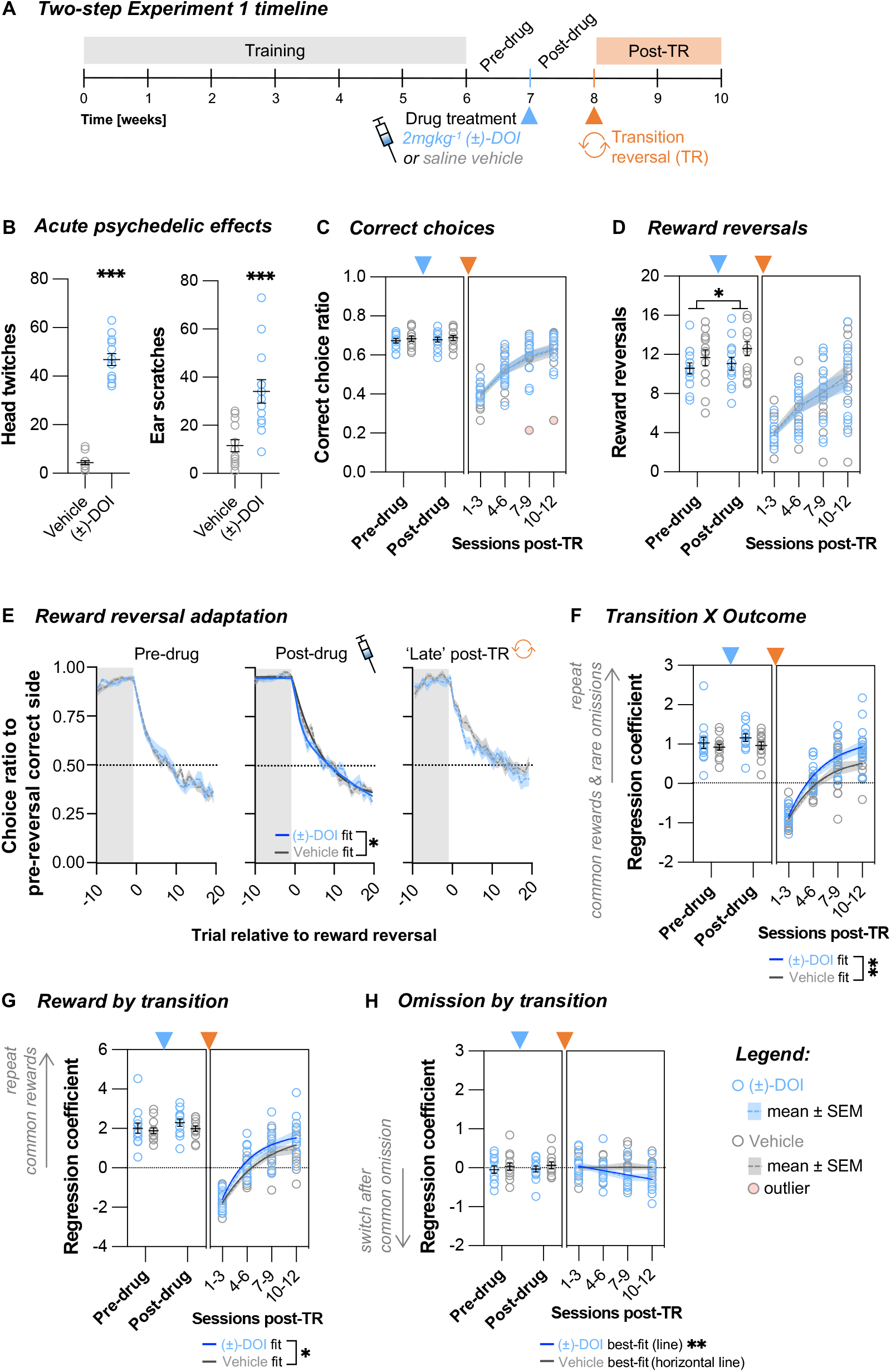
(±)-DOI administered one week before a novel transition reversal improved learning via a new choice strategy. **(A)** Two-step Experiment 1 timeline. After the animals were fully trained and had one week of stable performance on the two-step task (*Pre-drug* period), they were treated with either saline vehicle or 2mgkg^−1^ (±)-DOI. The day after treatment, testing was continued on the same version of the task for one week (*Post-drug* period). A single novel transition reversal (TR) was then initiated at the start of the second post-drug week. Animals’ performance was tracked for a further two weeks (*Post-TR* period). **(B)** (±)-DOI induced a high frequency head-twitch response (unpaired Welch’s *t*-test, *t*_14.9_=16.58, *P*<0.001, Cohen’s *d*=6.5, BF_10_>>100) and ear-scratch response (unpaired Welch’s *t*-test, *t*_17.8_=4.11, *P*<0.001, Cohen’s *d*=1.6, BF_10_=65.2). **(C)** The number of correct choices per session did not differ across treatment groups pre- or post-drug (two-way RM ANOVA: Time *F*_1,24_=0.42, *P*=0.521, BF_incl_=0.33; Drug *F*_1,24_=0.34, *P*=0.566, BF_incl_=0.50; Time X Drug *F*_1,24_=0.002, *P*=0.967, BF_incl_=0.36). Post-TR adaptation was not influenced by the drug (nonlinear one-phase association regression fits, *F*_3,96_=0.63, *P*=0.599, AICc_10_=0.09). **(D)** The number of reward reversals per session did not differ pre-drug, but it increased post-drug irrespective of the treatment type (two-way RM ANOVA: Time *F*_1,24_=4.3, *P*=0.048, η_p_^2^=0.15, BF_incl_=1.51; Drug *F*_1,24_=2.1, *P*=0.160, BF_incl_=0.95; Time X Drug *F*_1,24_=0.46, *P*=0.503, BF_incl_=0.41). Post-TR, the treatment groups remained comparable (nonlinear line regression fits, *F*_2,100_= 0.110, *P*=0.896, AICc_10_=0.13, slope_global_=1.9, CI [1.4, 2.3]). **(E)** The pre-drug reward reversal performance was comparable across treatment groups (double exponential fit permutation test, tau_fast_ *P*=0.671, tau_slow_ *P*=0.851, tau_fast/slow mix_ *P*=0.900). Post-drug, (±)-DOI-treated mice were quicker to reverse their choices (tau_fast_ *P*=0.070, tau_slow_ *P*=0.036, tau_fast/slow mix_ *P*=0.046). In the late post-TR period (post-TR sessions 7-12, when the animals restarted doing many reward reversals per session), the trend for faster adaptation in (±)-DOI-treated mice persisted, but the difference was not statistically significant (tau_fast_ *P*=0.130, tau_slow_ *P*=0.085, tau_fast/slow mix_ *P*=0.152). **(F)** Trial-to-trial learning of reward and transition probabilities was not different pre- and post-drug (two-way RM ANOVA: Time *F*_1,24_=1.4, *P*=0.246, BF_incl_=0.49; Drug *F*_1,24_=1.3, *P*=0.261, BF_incl_=0.66; Time X Drug *F*_1,24_=0.4, *P*=0.528, BF_incl_=0.41). (±)-DOI-treated mice were faster during the TR adaptation (nonlinear one-phase association regression fits, *F*_3,98_=4.76, *P*=0.004, AICc_10_=40.10). **(G)** Trial-to-trial reward learning was not different pre- and post-drug (two-way RM ANOVA: Time *F*_1,24_=3.4, *P*=0.079, BF_incl_=1.04; Drug *F*_1,24_=0.8, *P*=0.387, BF_incl_=0.61; Time X Drug *F*_1,24_=0.6, *P*=0.444, BF_incl_=0.44). (±)-DOI-treated mice were faster during the TR adaptation (nonlinear one-phase association regression fits, *F*_3,98_=2.82, *P*=0.043, AICc_10_=2.53). **(H)** Trial-to-trial reward omission learning was comparable pre- and post-drug (two-way RM ANOVA: Time *F*_1,24_=0.13, *P*=0.725, BF_incl_=0.29; Drug *F*_1,24_=0.76, *P*=0.392, BF_incl_=0.47; Time X Drug *F*_1,24_=0.01, *P*=0.907, BF_incl_=0.36). Post-TR, the best-fit of (±)-DOI-treated mice (horizontal line vs. line fit *P*=0.007, AICc_10_=14.81, slope_(±)-DOI_=-0.112, CI [−0.192, - 0.032]) was different to that of vehicle-treated mice (horizontal line vs. line fit *P*=0.868, AICc_10_=0.33, mean_Veh_=0.010, CI [−0.066, 0.086]). Trial-to-trial reward omission learning was not significant for either treatment group pre-TR (one sample *t*-tests, all *P*>0.452, BF_10_<0.36). Post-TR, vehicle-treated mice remained uninformed by omissions (one sample *t*-tests, all *P*>0.739, BF_10_<0.29), but (±)-DOI-treated mice started incorporating omissions in their choice strategy by the second post-TR week (one sample *t*-tests: [1–3] *t*_12_=0.87, *P=*0.403, BF_10_=0.38; [4–6] *t*_12_=-0.87, *P=*0.406, BF_10_=0.38; [7–9] *t*_12_=-2.82, *P=*0.015, BF_10_=4.08; [10–12] *t*_12_=-2.48, *P=*0.029, BF_10_=2.46). Data shown as mean ±SEM. *n*_Veh_=13. *n*_(±)-DOI_=13. \**P*<0.05. \*\**P*<0.01.

We next assessed how (±)-DOI affected choice performance specifically around the reward reversals. The speed of adaptation was comparable across all animals before drug treatment (Fig.3E left, *P*>0.671). However, (±)-DOI-treated mice exhibited a faster switch in their choice probability trajectory than the controls (Fig.3E centre, tau_fast_ *P*=0.070, tau_slow_ *P*=0.036, tau_fast/slow mix_ *P*=0.046). To examine the basis of this effect, we directly compared within-subject choice trajectories pre- and post-drug. This highlighted that the aforementioned effect was driven by the vehicle-treated mice being slower at reward reversals post-drug (tau_fast_ *P*=0.059, tau_slow_ *P*=0.018, tau_fast/slow mix_ *P*=0.097), whereas the (±)-DOI group’s performance, while marginally faster (tau_fast_ pre-drug 3.17 versus post-drug 1.28; tau_slow_ pre-drug 33.86 versus post-drug 24.79), was not statistically different post-drug (*P*>0.088).

Next, to determine if (±)-DOI affects adaptation to previously unencountered challenges, we initiated a single novel reversal in the transition structure at the end of the first post-drug week (Fig.2B, *transition reversal*, TR). As expected, there was a notable disruption in general task performance. Trial numbers remained intact (Fig.S5A), but mice initially made fewer correct choices (Fig.3C) and completed fewer reward reversals per session (Fig.3D). The subsequent recovery of these measures to pre-drug performance levels over time was not influenced by (±)-DOI (Fig.3C *P*=0.804, Fig.3D *P*=0.896). (±)-DOI-treated mice still appeared quicker at reward reversal adaptation in the second post-TR week (Fig.3E right), but the post-TR drop in general task performance confounded these measures, so the difference was no longer statistically significant (all *P*>0.091).

To measure cognitive adaptation to the TR, we tracked the animals’ trial-by-trial choice strategies across the two post-TR weeks, as measured by the logistic regression model (see Fig.2E). While the *Outcome* and *Transition* predictors quantify the simple reinforcing effects of outcome and transition type (Fig.S5B-C), the significantly positive interaction predictor, *Transition X Outcome* (Fig.3F), reflects the animal’s comprehension of the task’s probabilistic structure wherein transitions and outcomes are interconnected. In the early post-reversal sessions, the significantly negative *Transition* and *Transition X Outcome* predictors signal that the animals were initially following the original transition structure. As animals learned the new structure, predictor loadings shifted back towards positive values, allowing us to track animals’ evolving understanding of the task structure over time. (±)-DOI significantly influenced the *Transition X Outcome* coefficient levels (Fig.3F, *P*=0.004), illustrated by the (±)-DOI group’s higher rate constant and plateau (*P*=0.006), but not initial post-TR values (*P*=0.919). Therefore, despite the equal disruption by TR, (±)-DOI-treated mice exhibited a quicker strategy reversal, indicated by a higher influence of the *Transition X Outcome* interaction on subsequent choices throughout the adaptation period.

To explore factors contributing to the faster strategy reversal in (±)-DOI-treated mice, we examined potential changes in their learning strategy after the transition reversal by evaluating reward and omission learning separately. Post-TR trial-by-trial reward learning was significantly different (Fig.3G *P*=0.043) with (±)-DOI-treated mice consistently showing a higher tendency than the controls to repeat common rewards. In line with our pre-treatment performance analyses (see Fig.2G), both treatment groups had non-significant *Omission by transition* predictors pre-TR (Fig.3H, all *P*>0.452). Post-TR, vehicle-treated animals continued to exhibit consistent absence of learning from omissions (Fig.3H, best-fit: horizontal line model; all *P*>0.739). Strikingly, by contrast, the *Omission by transition* predictor of (±)-DOI-treated mice *decreased* over time (best-fit: line model preferred, *P*=0.007, slope −0.112, CI [−0.192, −0.032]), becoming significantly negative by the second week post-TR (*P*<0.029 for sessions 7-12). The negative loading reveals that (±)-DOI-treated animals started to be influenced by omissions and would be more likely to switch to a different choice after experiencing a common reward omission – an optimal strategy previously overlooked by the mice. We confirmed this effect was consistent across individual subjects, with 9 out of 13 (±)-DOI-treated mice exhibiting a more negative *Omission by transition* predictor after TR, compared to only 4 out of 13 vehicle-treated mice (Fig.S6, likelihood ratio test *P*=0.047).

To assess how unusual this emergence of omission sensitivity in (±)-DOI-treated mice is in the two-step task, we reanalysed the dataset from a previous study [54] using our logistic regression model that evaluates rewards and omissions separately. This showed a non-significant *Omission by transition* predictor, consistent with our current experiments (Fig.S7A). We then simulated pre-drug behaviour using an asymmetric inference model drawn from the distribution of fits from all *N*=82 animals tested on the two-step task to again confirm that choice strategies were only influenced by rewarded but not unrewarded trials (Fig.S7B left). By contrast, the post-TR behaviour of (±)-DOI-treated animals in Two-step Experiment 1 exhibited a significant influence of omission trials in their choice strategy (Fig.S7C), now exemplifying symmetric inference learning (Fig.S7B right). Note, however, that although (±)-DOI-treated animals started using information from both rewards and omissions, they did not give equal value to both types of outcomes, with rewarded outcomes still having greater impact on stay/switch behaviour.

We then repeated our initial *ex vivo* MRI analyses for a subset of brains collected at the end of Two-step Experiment 1 (*n*_group_=10) to look at possible brain structural differences between (±)-DOI- and vehicle-treated mice. However, at this timepoint, we found no significant differences in grey-matter volume in either the voxel-wise or ROI-wise analyses (linear model drug effects not significant at *q*>0.20, Fig.S8), even when we used seeded analysis with the ROIs that had exhibited significant differences one day post-treatment in our first imaging study (Fig.S9).

Taken together, these data indicate that (±)-DOI did not enhance overall decision-making accuracy but resulted in a faster change in strategy during a novel transition reversal. Crucially, (±)-DOI-treated mice exhibited a unique shift in strategy and started learning from reward omissions, which was not observed in the vehicle-treated mice.

### The importance of post-drug timing

Our findings in Two-step Experiment 1 suggested that there might be a temporal dissociation between measurable brain structural changes and the observed cognitive enhancements. To investigate the relationship between the timing of drug administration and cognitive flexibility, in a second cohort (Fig.4A-B, *Two-step Experiment 2*, *N*=29), we initiated the transition reversal one day after drug treatment. This is when previous studies have described neuronal plasticity enhancements [17–21] and when our initial MRI study showed significant structural plasticity changes (Fig.1). There were two possible scenarios: either (±)-DOI’s effects on novel reversal learning are correlated with increases in neuronal plasticity, suggesting *stronger* effects when testing adaptability to a novel TR sooner after drug treatment; or there is a critical time component whereby a period of strengthening and integration of newly formed synaptic connections is necessary to utilize the burst of neuronal psychedelic-induced plasticity in imparting long-lasting cognitive effects, suggesting *weaker* effects if testing novel reversal learning immediately after drug treatment.

**Fig.4.**
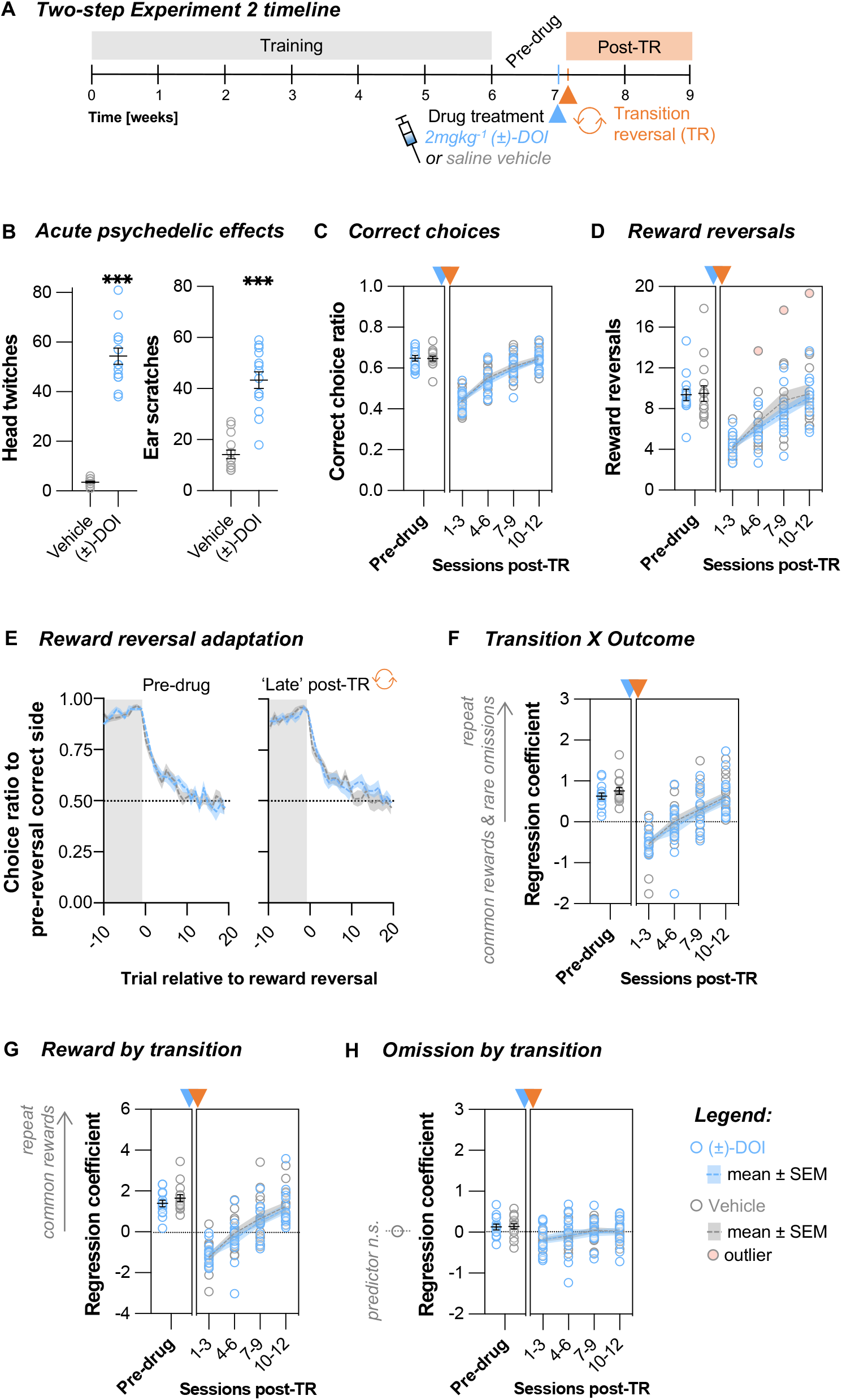
(±)-DOI administered one day before a novel transition reversal did not lead to improved learning. **(A)** Two-step Experiment 2 timeline. The animals were treated with either saline vehicle or 2mgkg^−1^ (±)-DOI after they were fully trained and had one week of stable performance on the two-step task (*Pre-drug* period). A transition reversal (TR) was initiated the day after drug treatment. Animals’ performance was tracked for a further two weeks (*Post-TR* period). **(B)** (±)-DOI induced a high frequency head-twitch response (unpaired Welch’s *t*-test, *t*_13.3_=15.61, *P*<0.001, Cohen’s *d*=5.9, BF_10_>>100) and ear-scratch response (unpaired Welch’s *t*-test, *t*_19.5_=7.96, *P*<0.001, Cohen’s *d*=3.0, BF_10_>>100). **(C)** The number of correct choices per session was not significantly different across treatment groups pre-drug (unpaired *t*-test, *t*_27_=0.096, *P=*0.924, BF_10_=0.35) or post-TR (nonlinear line regression fits, *F*_2,112_=0.22, *P*=0.804, AICc_10_=0.14, slope_global_=0.067, CI [0.057, 0.076]). **(D)** The number of reward reversals per session was not different pre-drug (Mann-Whitney test, W=86.5, *P=*0.431, BF_10_=0.42) or post-TR (nonlinear line regression fits, *F*_2,109_=0.00008, *P*>0.999, AICc_10_=0.11, slope_global_=1.6, CI [1.3, 1.9]). **(E)** The pre-drug reward reversal performance was not different across treatment groups (double exponential fit permutation test, tau_fast_ *P*=0.102, tau_slow_ *P*=0.100, tau_fast/slow mix_ *P*=0.244). In the late post-TR period, the two treatment groups remained comparable (tau_fast_ *P*=0.169, tau_slow_ *P*=0.520, tau_fast/slow mix_ *P*=0.508). **(F)** Trial-to-trial learning of reward and transition probabilities was not different pre-drug (Mann-Whitney test, W=120, *P=*0.533, BF_10_=0.46) or post-TR (nonlinear line regression fits, *F*_2,111_=0.17, *P*=0.837, AICc_10_=0.14, slope_global_=0.35, CI [0.28, 0.43]). **(G)** Trial-to-trial reward learning was not significantly different pre-drug (Mann-Whitney test, W=123, *P=*0.451, BF_10_=0.48) or post-TR (nonlinear line regression fits, *F*_2,112_=0.30, *P*=0.738, AICc_10_=0.16, slope_global_=0.79, CI [0.65, 0.93]). **(H)** Trial-to-trial reward omission learning was not significantly different pre-drug (unpaired *t*-test *t*_27_=0.15, *P=*0.880, BF_10_=0.35) or post-TR (nonlinear horizontal line regression fits, *F*_1,114_=0.033, *P*=0.856, AICc_10_=0.35, mean_global_= −0.055, CI [−0.116, 0.006]). Data shown as mean ±SEM. *n*_Veh_=15. *n*_(±)-DOI_=14.

The post-TR recovery of general task performance was again not affected by (±)-DOI. Task engagement (Fig.S5D), choice accuracy (Fig.4C), the number of reward reversals (Fig.4D) and speed of reward reversal adaptation (Fig.4E) were not significantly different across treatment groups post-TR (all *P*>0.508). However, in contrast to Two-step Experiment 1, the rate at which the animals learned the new transition structure was comparable across treatment groups. The *Transition X Outcome* (Fig.4F, *P*=0.837) and *Reward by transition* (Fig.4G, *P*=0.738) predictors were equivalent in both (±)-DOI and vehicle mice, and, crucially, learning from omissions remained negligible in both groups (Fig.4H, *P*=0.856).

These findings imply that (±)-DOI’s effects on cognitive flexibility were not immediate, emphasizing a critical time component in shaping how (±)-DOI influences cognitive flexibility and the development of novel behavioural strategies.

### The importance of post-drug experience

The timing of drug administration relative to TR is not the only factor that was different between Two-step Experiments 1 and 2. In the first cohort, during the first post-drug week, before TR, the mice continued accumulating task experience. Marginal increases in the number of completed trials (Fig.S5A) and reward reversals (Fig.3D) were observed, regardless of the treatment they received. Hence, the observed cognitive effects may have been influenced not just by the timing of TR, but by the experiential context available during that time too. Indeed, psychedelic drug research has continuously highlighted the importance of psychological and environmental factors in shaping the behavioural effects of psychedelic drugs [23, 55].

Therefore, to investigate the influence of training in mediating (±)-DOI’s effects on cognitive flexibility, in our final cohort of animals, we deliberately prevented any further task experience during the first post-drug week (Fig.5A-B, *Two-step Experiment 3*, *N*=27). Our prediction was that if the training on the original task during the post-drug period facilitated the necessary experience for cognitive plasticity processes, then the absence of this experiential component would attenuate the reversal learning effects observed.

**Fig.5.**
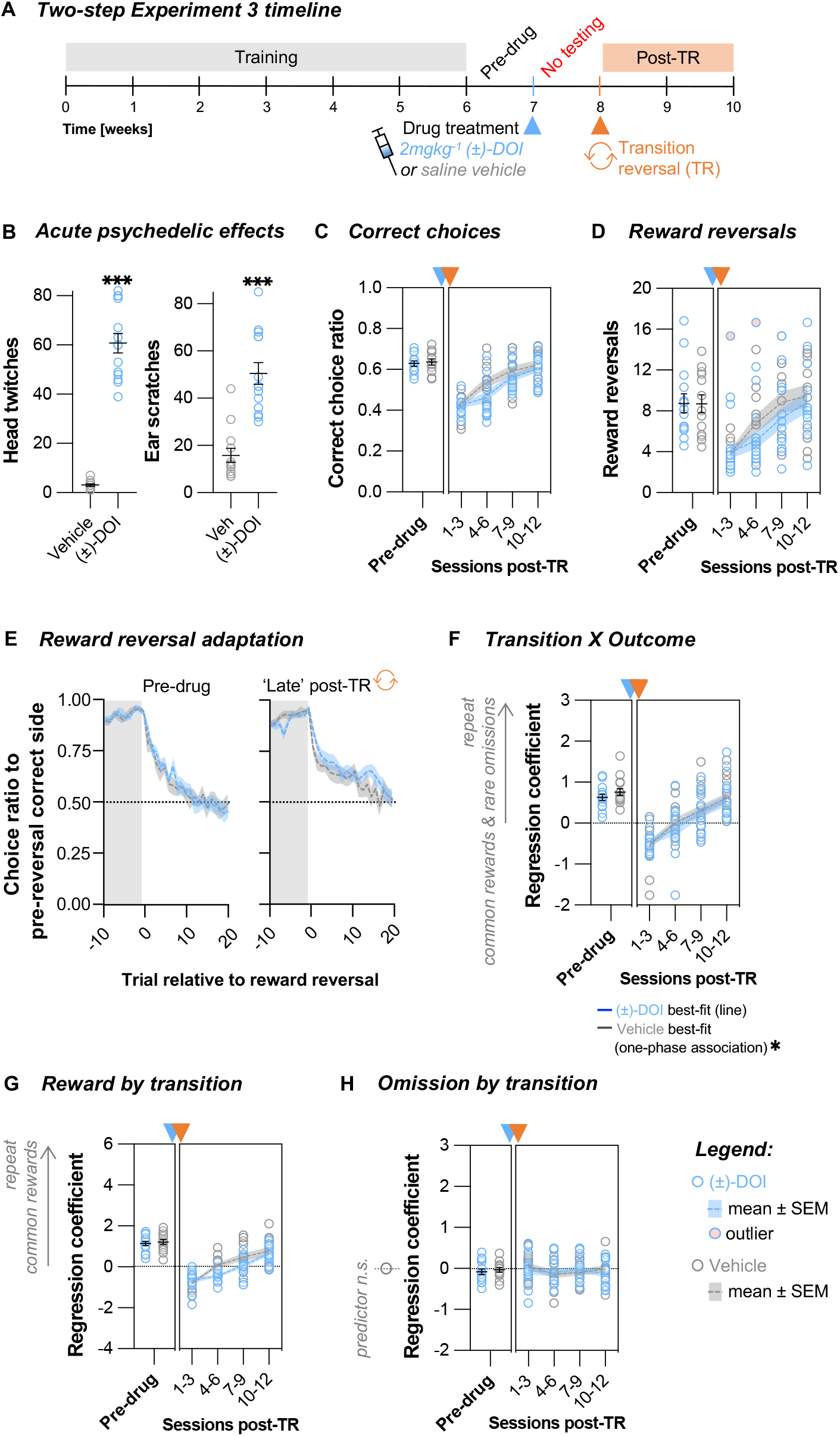
Effects of (±)-DOI on the adaptability to a novel reversal occurring one week after treatment were dependent on experience. **(A)** Two-step Experiment 3 timeline. The animals were treated with either saline vehicle or 2mgkg^−1^ (±)-DOI when they were fully trained and had one week of stable performance on the two-step task (*Pre-drug* period). For the next week, the animals were kept under water deprivation but were not allowed any further experience on the two-step task. A novel transition reversal (TR) was then initiated at the start of the second post-drug week. Animals’ performance was tracked for a further two weeks (*Post-TR* period). **(B)** (±)-DOI induced a high frequency head-twitch response (unpaired Welch’s *t*-test, *t*_13.4_=14.24, *P*<0.001, Cohen’s *d*=5.5, BF_10_>>100) and ear-scratch response (Mann-Whitney U test, W=7, *P*<0.001, Rank-Biserial Corr.=0.92, BF_10_=28.8). **(C)** The number of correct choices per session did not differ across treatment groups pre-drug (unpaired *t*-test, *t*_25_= −0.381, *P=*0.707, BF_10_=0.38) or post-TR (nonlinear line regression fits, *F*_2,104_=2.24, *P*=0.111, AICc_10_=1.11, slope_global_=0.064, CI [0.051, 0.077]). **(D)** The number of reward reversals per session was not significantly different pre-drug (unpaired *t*-test, *t*_25_= −0.045, *P=*0.965, BF_10_=0.36) or post-TR (nonlinear line regression fits, *F*_2,102_=2.54, *P*=0.084, AICc_10_=1.48, slope_global_=1.7, CI [1.2, 2.2]). **(E)** The pre-drug (double exponential fit permutation test, tau_fast_ *P*=0.513, tau_slow_ *P*=0.354, tau_fast/slow mix_ *P*=0.610) and late post-TR (tau_fast_ *P*=0.560, tau_slow_ *P*=0.719, tau_fast/slow mix_ *P*=0.368) reward reversal performance was comparable across treatment groups **(F)** Trial-to-trial learning of reward and transition probabilities was not different pre-drug (unpaired *t*-test *t*_25_=0.123, *P=*0.903, BF_10_=0.36). (±)-DOI group’s fit was significantly different post-TR. Nonlinear regression fits: (±*)-DOI*, line fit (one-phase association model failed to converge on a best-fit curve), slope=0.064, CI [0.051, 0.077]; *Vehicle*, line vs. one-phase association fit: *F*_1,49_=6.19, *P*=0.016, AICc_10_=6.79. **(G)** Trial-to-trial reward learning was not different pre-drug (unpaired *t*-test, *t*_25_=0.449, *P=*0.657, BF_10_=0.39) or post-TR (nonlinear line regression fits, *F*_2,104_=3.01, *P*=0.054, AICc_10_=2.36, slope_global_=0.47, CI [0.39, 0.56]). **(H)** Trial-to-trial reward omission learning was not significantly different pre-drug (unpaired *t*-test, *t*_25_=0.563, *P=*0.578, BF_10_=0.40) or post-TR (nonlinear horizontal line regression fits, *F*_1,106_=0.229, *P*=0.633, AICc_10_=0.39, mean_global_= −0.064, CI [−0.125, −0.002]). Trial-to-trial reward omission learning was not contributing to the choice strategy of either treatment group (one sample *t*-tests, all *P*>0.095, BF_10_<0.97). Data shown as mean ±SEM.. *n*_Veh_=13. *n*_(±)-DOI_=14.

In contrast to previous cohorts, there was a noticeable decline in task engagement in both treatment groups following the off-task week, but trial rates rebounded upon resuming daily testing after the transition reversal (Fig.S5G) and the extent of this effect was not different across treatment groups (*P*=0.306). Consistent with Two-step Experiments 1 and 2, the post-TR recovery of general task performance, including task accuracy (Fig.5C), reward reversal rate (Fig.5D), and speed of adaptation (Fig.5E), did not show significant differences between the treatment groups (all *P*>0.084). Nonetheless, a trend was visible suggesting that (±)-DOI-treated animals now exhibited a marginally slower recovery compared to the control group (Fig.5C-D).

Examining the cognitive adaptation more closely, specifically looking at the *Transition X Outcome* predictor, we observed a significant effect of (±)-DOI, but, contrary to Two-step Experiment 1, (±)-DOI-treated mice exhibited a *slower* rather than a faster rate of adaptation than the vehicle group (Fig.5F, vehicle best-fit one-phase association preferred over line regression, *P*=0.016). While differences in the *Transition* (Fig.S5I, *P*=0.055) and *Reward by transition* predictor (Fig.5G, *P*=0.054) did not quite reach statistical significance, the trend for (±)-DOI-treated mice being slower than the controls to change was still present. Moreover, there was no evidence for learning from omissions in either group (Fig.5H, all *P*>0.095), so the faster adaptation in vehicle-treated animals was not accompanied by a change in strategy, unlike the one observed in (±)-DOI-treated mice in Two-step Experiment 1. Therefore, the absence of additional task experience during the post-drug week not only curtailed the cognitive changes observed in Two-step Experiment 1 but also resulted in the opposite direction of effect.

Together, our three separate two-step experiments underscored the importance of both time- and experience-dependent mechanisms in influencing (±)-DOI’s effects on the speed of novel reversal learning. To assess whether the changes observed in *Transition X Outcome* predictor differed significantly across our two-step experiments, we conducted a 3-way ANOVA with the within-subjects factor of post-TR session, and between-subjects factors of drug and experiment (Session *P*<0.001, η_p_^2^=0.70, Session X Experiment *P*<0.001, η_p_^2^=0.21, Drug X Experiment *P*=0.035, η_p_^2^=0.08, rest *P*>0.176). Crucially, we found a significant drug by experiment interaction, confirming that the effects of (±)-DOI on cognitive flexibility varied across the two-step experiments, highlighting the importance of considering the experimental context and timing when interpreting the drug’s effects.

## Discussion

Here we demonstrate how (±)-DOI treatment facilitated a faster change in strategy during a novel transition reversal without affecting overall decision-making accuracy. Remarkably, (±)-DOI was able to induce a unique shift in the learning strategy whereby mice started learning from previously overlooked reward omissions. Crucially, we further uncovered the complex time- and context-dependent nature of (±)-DOI’s effects on cognitive flexibility. The timing of drug administration and the availability of post-drug training were both found to be critical, as cognitive effects were not immediate and depended on post-drug task experience.

We show that, in the first week post-treatment, (±)-DOI-treated animals were quicker than the controls to reverse their choices following serial reversals in reward probabilities. This difference was driven, in part, by an apparent post-treatment drop in performance in control animals. This could indicate that (±)-DOI’s effect on serial reward reversal adaptation may be that of improving cognitive stability [56] – a protective effect of maintaining optimal cognitive performance against a drop seen in the control group and possibly induced by injection stress. Alternatively, (±)-DOI might promote serial reversal learning to rescue a deficit, but further enhancements were limited by a putative ceiling effect due to the extensive training on such reversals. Recent data show that tabernanthalog, a plasticity-inducing non-psychedelic analogue of the psychedelic drug ibogaine, was able to rescue a serial reversal learning deficit in mice that underwent unpredictable mild stress [57]. Tabernanthalog, like (±)-DOI in our study, did *not* increase performance past the levels seen in the non-treated controls, suggesting no added performance benefit due to the drug.

(±)-DOI’s ability to shape cognitive flexibility was further revealed when the animals were tested on a single novel reversal in the task’s transition structure, suggesting (±)-DOI’s effects could be distinctive if testing single versus serial reversal learning. When the transition reversal was implemented one week after drug treatment, (±)-DOI-treated animals were better able to inhibit responding to the old model. Strikingly, they also now incorporated a novel learning strategy whereby they exhibited significant sensitivity to reward omissions. This is notable as omissions were otherwise disregarded by mice, a pattern replicated in previous experiments using the two-step [54] and other decision-making tasks [58–60]. One intriguing possibility is that the added challenge of a novel kind of reversal and the frustrative increased rate of reward omissions may have acted as an acquired motivation to change strategies [61], which could have primed (±)-DOI-mediated increased sensitivity to the previously overlooked omission cues. Notably, learning asymmetry for reward versus omission is not unique to mice. Humans solving the two-step task and other reward-guided probabilistic tasks also show higher learning rates for positive feedback [62–65], and reduced loss aversion has been implicated in addictive disorders such as gambling and alcohol dependence [66]. Our finding that (±)-DOI aids faster learning from both rewards and omissions echoes the recent human data suggesting LSD increased both reward *and* punishment learning rates in healthy subjects doing a probabilistic reversal task, although the human subjects in this study were performing the task in the acute phase of drug action [67].

The cognition enhancing effects of psychedelics are widely believed to result from the enhancement of neural plasticity processes [2] which are presumed to be ramping up over the first week following treatment [18, 19, 21], so the time between drug treatment and any novel reversal are potentially critical for the observed effects. In fact, we saw no adaptability differences with (±)-DOI when we initiated the transition reversal one day after drug treatment, suggesting that while some cognitive benefits may develop during the sub-acute phase, as we saw some benefit of (±)-DOI on reward reversals in the first post-drug week, but additional time may be needed to confer benefits for novel adaptations to more challenging task changes. One possibility is that the consolidation of cellular plasticity changes is a predecessor for, and a modulator of, lasting improvements in cognitive flexibility. “Consolidation” here refers to the process of synaptic pruning and strengthening that determines which spines are retained and integrated into neural networks in the longer-term. How the pruning process of psychedelic-induced neuronal plasticity is regulated, and whether it can be directed for a specific purpose, is an intriguing area of future research.

Remarkably, we also found that (±)-DOI did not confer cognitive benefits when we did not allow the animals further task experience after drug treatment, before then reversing the task structure. Instead, (±)-DOI had a mild detrimental effect, slowing down the animals’ relearning. One explanation for this could be the lack of enriched behavioural experience during the critical neuroplasticity window. Barring the animals from the task restricted their cognitive stimulation, which is known to be a positive modulator in the environment capable of inducing neuronal plasticity [68]. Reduced cognitive activity could be perceived as environmental impoverishment [69], whose detrimental effects on learning and/or downscaling of plasticity processes could be exacerbated by (±)-DOI. It has been suggested that neuroplasticity induced by serotonin-enhancing drugs acts as a catalyst for a “unidirectional susceptibility to change” that allows subsequent stimuli to reshape neural circuits in both positive and negative directions, depending on the quality of the environment [22, 70]. Neuroplasticity is a complex sequence of molecular, cellular, and network-level events that unfold across time, and, at each level, experience may modulate the course of this intricate cascade, for example, by directing the consolidation of synaptic remodelling induced by psychedelic drugs.

Consistent with the widely replicated neuronal plasticity-promoting effects of (±)-DOI [17, 19, 20] we also provide the first evidence that a single moderate dose of (±)-DOI can induce regional volume changes across sensory and association cortical areas, as well as subcortical structures, beyond the acute phase of drug action. These discoveries complement previous findings of psychedelic-induced increases in dendritic branching and spine formation [17–21, 25, 26], and suggest that (±)-DOI may have widespread effects on whole-brain structural plasticity. Notably, our effects in S1 complement previous reports of (±)-DOI-induced spinogenesis in that region [19], and acute changes in activity measured in pharmacological MRI studies with psychedelics [71, 72]. We did not observe any significant volume changes three weeks post-treatment. This is consistent with prior evidence suggesting that the rates of structural plasticity changes are at their highest in the first few days after psychedelic drug treatment [19–21] and that only a third of newly formed spines are maintained three weeks after [18]. Therefore, any remaining differences in spine density may be too subtle to be detected with a whole-brain analysis. Furthermore, long-term cognitive and behavioural changes may not require all of the initial growth of spines and dendritic branches to be maintained across time. The initial neuroplasticity burst could instead be acting as the foundation for rewiring circuits and changing network dynamics which, in turn, then directs changes in higher-level processes, such as cognitive flexibility.

Our findings highlight the multifaceted nature of psychedelics’ therapeutic effects and their relevance for various mental health conditions. We suggest that these therapeutic benefits may not be solely dependent on the initiation of neuroplasticity but could also be influenced by the specific post-treatment experiential context. These insights prompt further mechanistic studies of how the pharmacological plasticity-promoting effects of psychedelics can be synergistically combined with training-guided network reorganization, leveraging the increased plasticity induced by the drug. Such an integrated approach holds promise for enhancing the therapeutic potential of psychedelics, particularly for patients exhibiting perseverative behaviours and a resistance to change, which is commonly observed in depression, anxiety, and addictions. The ability of psychedelics to both facilitate adaptive responses and incorporate previously unexplored information may bring new opportunities for promoting positive therapeutic outcomes in clinical settings where rigid cognitive patterns and resistance to traditional interventions pose a particular challenge.

## Supplementary information

Supplementary information includes Supplementary Methods, Supplementary Tables 1 and 2, and Supplementary Figures 1-9.

## Acknowledgements

We thank Claire Bratley, Lily Qui, Mohamed Tachrount, Urs Schuffelgen, Clemence Ligneul, Antoine Cherix, and Cristiana Tisca for technical assistance with *ex vivo* MRI sample and image collection. We also thank Thomas Akam for technical assistance with the two-step task and inputs on the data analysis, Vladyslav Vyazovskiy for useful discussions about the data, and Katie Hewitt for laboratory facility management and support. This research was funded in whole, or in part, by Wellcome (MŠ: 4-year studentship 102170/Z/13/Z; MEW: Senior Research Fellowship fellowship 202831/Z/16/Z and Collaborative Award 214314/Z/18/Z; JPL and the Wellcome Centre for Integrative Neuroimaging: supported by core funding from the Wellcome Trust 203139/Z/16/Z). For the purpose of open access, the author has applied a CC BY public copyright licence to any Author Accepted Manuscript version arising from this submission.

## Author contributions

MŠ: conceptualization, methodology, software, formal analysis, investigation, data curation, visualization, writing—original draft, review and editing, funding acquisition. AL: software, formal analysis, writing—review and editing. MB-P: methodology, software, formal analysis, writing—review and editing. JPL: methodology, resources, supervision, funding acquisition, writing—review and editing. DMB: conceptualization, methodology, resources, supervision, writing— review and editing. MEW: conceptualization, methodology, resources, supervision, funding acquisition, writing—review and editing.

## Conflict of interest

MEW and DMB have received research funding from COMPASS Pathways to start in 2023. However, the work described here was conducted prior to this agreement and was entirely independent of any input or influence from the company. The authors affirm that COMPASS Pathways had no involvement in the design, data collection, analysis, interpretation, or preparation of this manuscript. The authors have no other competing interests to declare.

