## Supplementary Information for "Lasting dynamic effects of the psychedelic 2,5-dimethoxy-4-iodoamphetamine ((±)-DOI) on cognitive flexibility"

### Supplementary Methods

##### *Animals*

C57BL/6J mice (Strain Code 632, Charles River, Kent, UK) were between 5- and 6-week-old at arrival, after which they were acclimatized for one week and ear punched for identification. The housing room was temperature-(21±2°C) and humidity-controlled (55±10%) on a 12h light/dark cycle (lights off at 7PM), with all procedures done during the light phase. All animals were housed in groups of 3-4 in open-top cages with sawdust and enrichment available (sizzle nest, and cardboard tunnel and house). All cage mates underwent procedures at the same time. We minimized variability in behavioural experiments by: (i) testing animals of a similar age, (ii) testing at approximately the same time of day throughout the experiment, (iii) using the same experimental protocol for two-step experiments across different cohorts, and (iv) using a standardized way of tunnel handling to minimize stress [1]. Numbers of animals used for experiments are stated in the figure legends. We aimed for a minimum of 10-15 animals per condition for our behavioural experiments based on our prior studies [2, 3].

##### *Water restriction*

Mice tested in the two-step task were water-restricted for motivation. At 6 weeks old, animals were weighed for two consecutive days to give an averaged pre-restriction baseline bodyweight for each animal. This baseline was age-corrected by 2% every week until 14-weeks-old, after which a 1%

correction was used. One day after starting water restriction, the animals were given 1h of unrestricted water access in their home cage, and behavioural training commenced the next day. During all training and testing sessions, the animals were weighed before and after the task, and if their bodyweight dropped below 85%, they had free water access in their home cage with food available until the bodyweight percentage was above 85%. All animals had a food pellet available in the operant box during behavioural training and testing. Animals were given 1h of free water access in their home cage once a week when they did not undergo two-step task testing. On the day of drug administration, injections were performed only after the two-step task testing was completed for the day, such that the animals already had their daily water access. In Two-step Experiment 3, during the first week after drug treatment, when the animals were barred from task training and could not get their water during the task, the mice were allowed a maximum 5min of free water access during which they consumed 1-2ml of water. In this way, the mice were kept water deprived throughout the experiment while minimizing the bodyweight increase in this period of more frequent free water access.

***Drug***

Racemic 2,5-dimethoxy-4-iodoamphetamine ((±)-DOI) is a synthetic phenethylamine psychedelic that is historically the most extensively used psychedelic in rodent preclinical research [4]. Its use in humans is limited due its long duration of effects, although the quality of the acute experience has been reported as similar to LSD [5]. (±)-DOI is not only more commercially available for use in research, but it also has well-characterized receptor binding and activation profiles, and behavioural effects in animals [4, 6–8]. (±)-DOI is highly soluble and chemically stable in solution and is one of the few psychedelics considered to be *relatively* selective. (±)-DOI is a potent agonist for most serotonin type 2 receptors (5-$HT_2$Rs), but it is reported to have up to 40-fold higher affinity for 5-$HT_{2A}$R than for 5-$HT_{2C}$R sites [9]. The plasticity-promoting effects of (±)-DOI are comparable to LSD, DMT, and psilocin/psilocybin in terms of the increase in dendritic arborization and spine formation [10, 11]. Behavioural therapeutic-like effects have also been found across different types of psychedelics in animals [10–14] and in humans [15–21]. Therefore, since these psychedelics share the plasticity-promoting effects that are a likely common mechanism of any behavioural effects, we expect to find analogous cognitive and structural effects with other psychedelic drugs to the ones we report here with (±)-DOI, though their relative efficacy could differ according to their distinct pharmacological profiles.

To determine what dose of (±)-DOI to use for our experiments we performed an experiment in which we determined dose-response curves for (±)-DOI's acute psychedelic-like effects – the head-twitch [4] and ear-scratch response [22]. To assess these acute psychedelic effects of (±)-DOI, head-twitches and ear-scratches were video recorded (Revotech I706-POE camera, max. 20fps) in a 42×20×20cm clear Plexiglass box in the first 30min after injection and quantified manually by the same trained experimenter for all studies. The recognizable frequency of head-twitches and ear-scratches effectively unblinded the video analysis.

We tested three different doses of (±)-DOI: 0.5, 1, and 2mgkg$^{-1}$. Additionally, we observed the injected mice in two different environments, familiar or novel, since psychedelic effects are known to be context-dependent [23]. While previous reports in rodents mainly focused on physical stressors [24–26], we wondered if injecting the animals in a novel versus familiar environment would result in a different intensity of acute psychedelic effects. Therefore, to determine the optimal dose *and* environment for psychedelic treatment, we compared the dose-dependent acute psychedelic effects observed in a novel environment (the animals were not exposed to the recording box prior to injection) or a familiar environment (the animals were habituated to the recording box by allowing them 15min of daily free exploration for 5 consecutive days before injection, Fig.S1A). Each animal tested received only one injection (saline vehicle control, or one dose of (±)-DOI) in one of the environments (novel or familiar), according to a between-subjects design.

We confirmed that the animals were responding to the two environments differently on the day of injection by placing the animals in the recording box for 5min of free exploration before injection. We observed that the animals that had been habituated to the recording box travelled significantly lower distances than the animals who had only been placed in the cage for that observation period on that day (Fig.S1B). After injecting the animals, the numbers of head-twitches and ear-scratches were manually scored offline from recordings of the first 30min after injection. Dose-dependent increases in the number of head-twitches and ear-scratches were modulated by environmental novelty (Fig.S1C-D). Overall, the dose-response curves for (±)-DOI were in the novel environment suggested greater peak responses and peak responses at higher drug dose than in the familiar environment.

We wanted to use the dose that resulted in near-peak acute psychedelic effects, since higher doses would have a more mixed receptor signalling activation profile and could induce a serotonin syndrome, which we wanted to avoid. We chose the 2mgkg$^{-1}$ dose injected in a novel environment for our experiments since this was the dose that resulted in near peak head-twitch and ear-scratch responses.

**Ex vivo *magnetic resonance imaging (MRI)***

Structural brain plasticity measures are commonly limited to examining selected brain regions, either because of the technology (e.g., the limited field of views in multi-photon microscopy) or by the amount of labour required for whole-brain coverage (e.g., in immunohistochemistry). However, MRI enables us to rapidly search the whole brain for regions undergoing structural plasticity changes. Lab mice maintained in the same environment have similar brain anatomy and there is low natural variability of regional brain volumes [27], but MRI signals can be altered by changes in neuronal structure at the level of dendrites and synapses [28–30]. To assess sub-acute changes in regional grey matter volumes, we collected the brains of 9-week-old mice 24-36h after they had been injected with either 2mgkg$^{-1}$ (±)-DOI or saline vehicle. These animals were injected in either a familiar or a novel testing environment (Fig.S1), but the imaging study was not powered to test for any putative effect of environment. To assess long-term volume changes, we collected brains of 17-week-old mice after they completed testing in a behavioural reversal learning task, three weeks after 2mgkg$^{-1}$ (±)-DOI or saline injection. Based on published power analyses [31], by using ≥8 mice we expected to have 80% power for recovering volume differences of ≥10% at the explorative 20% false discovery rate (FDR) significance threshold.

*Sample collection*

Mice were anaesthetized via intraperitoneal 10mlkg$^{-1}$ injection of 150mgkg$^{-1}$ ketamine (Ketamidor, Chanelle Pharma) and 10mgkg$^{-1}$ xylazine (Rompun, Bayer Pharmaceuticals) in saline. Animals were perfused through the left ventricle, first with 30ml of phosphate-buffered saline (PBS), 1µlml$^{-1}$ of heparin (Wockhardt UK Ltd), and 2mM of Gadovist contrast (Bayer Pharmaceuticals) administered at a rate of 1ml per minute, and second with 30ml of 4% paraformaldehyde aqueous solution (PFA) and 2mM of Gadovist in PBS at the same rate. After perfusion, the mouse was decapitated and extracranial tissue removed, leaving the brain inside the skull. The skull was soaked in 4% PFA and 2mM Gadovist overnight at 4°C, then transferred to a solution of 2mM Gadovist in PBS and 0.02% sodium azide. The samples were stored at 4°C for a minimum of one month before MRI acquisition.

Pre-scanning sample preparation involved first placing the samples in a vacuum pump for 30-60min to remove any intracranial air-bubbles formed during storage. Any residual PBS was removed, and the sample was transferred to a holder filled with Fluorinert, a proton-free solution which minimizes MRI susceptibility artefacts. Samples were then kept at room temperature for 24h before scanning and during the acquisition. Following sample positioning in the scanner, a short multi-gradient echo (MGE) sequence was acquired to check for any remaining air-bubble artefacts. MGE sequence parameters were: 20 echoes, TE=3ms, 3ms inter-echo spacing, 200μm isotropic voxel resolution, and 120 X 40 X 55 matrix size.

*Data processing and analysis*

After quality control and pre-processing, we used the Mouse Build Model pipeline from Pydpiper [32] for non-linear deformation of aligning individual samples to the registered study average and extracting Jacobian determinants (JDs). Region of interest (ROI) segmentations were done based on the DSURQE atlas [33] through the MAGET algorithm [34]. ROI-wise analysis was unseeded and based on the hierarchical tree using ontogeny from the Allen Brain Institute to compute JDs for each structure in each hemisphere separately. Multiple comparisons were controlled with Benjamini and Hochberg method or Benjamin and Yekutieli method for voxel-wise and ROI-wise comparisons, respectively. The resulting maps were visualized on the study template for the voxel-wise analysis, or the DSURQE atlas for the ROI-wise analysis.

##### ***Two-step reversal learning task***

In this task, choices are a function of reward and transition history. If animals did not have an understanding of the task structure, subjects would be expected to repeat the same step 1 choice if the previous trial was rewarded, regardless of whether the transition between the steps was common or rare. While this is computationally simple, it is not very flexible. Creating an internal representation of the task structure, to track both the reward history and the task's transition structure, confers flexibility, as future implications of new information can be evaluated using the learned representations and not by trial and error [35–37].

Fifteen custom-built 12 X 12cm operant boxes, controlled using pyControl [38], were used to run the two-step task. Five nose poke ports (Fig.2A) were located on one of the operant box walls. A central port was flanked by a step 1 choice ports 4.0cm to the left and right, and by step 2 state ports 1.6cm above and below the central poke. A mouse initiated a trial by poking a central port and then it either (i) chose between a left or right port (*free choice* trial) or (ii) selected the one left/right port which was lit up (*forced choice* trial, included to ensure mice continuously explore both step 1 options). The choice/selection triggered a transition to the step 2 state where only one of the up/down ports lit up, selection of which triggered a probabilistic delivery of water reward. The next trial was initiated after a random 2-4s inter-trial interval. The two step 2 ports had a solenoid for delivering water rewards. A speaker located above the ports delivered auditory stimuli. An active port, i.e., the port that a mouse could interact with by nose poking, was indicated by illuminating that port. To ensure mice knew when they had made a nose poke on the active port, a click sound was presented whenever the mice poked the illuminated port. No click sound was presented if a mouse was poking inactive ports.

Reward probabilities of step 2 states reversed serially in blocks which could be non-neutral (reward probabilities switch between 80% and 20%) or neutral (both reward probabilities 50%). Reversals from non-neutral blocks were based on the animals' performance. If the exponential moving average of correct choices (tau =8 free choices) >75%, a reversal was triggered following a random delay of 5-15 trials. Reversals from neutral blocks were triggered after a random 20-30 trials interval. Transition probabilities between the two steps, which could be Type A (left→up & right→down in 80% of trials, vice versa in the other 20%) or Type B (left→down & right→up in 80% of trials, vice versa in the other 20%) were fixed during training and counterbalanced across animals.

*Training*

Training consisted of multiple stages with increasing complexity to build the sequence of multiple steps required by the task (Table S2). Training and testing occurred as one session per day, for 6 days per week. The sessions in stages 1.1-4.6 lasted 60min, but 90min sessions were used at the final training stage and throughout subsequent testing. Water reward size in the task started at 15µl and decreased to 4µl across training to increase the number of trials and reward reversals that the animals were doing per session. Animals required 10-26 sessions to reach the final stage of training (the times taken for separate stages are indicated in Table S2).

At the training stage 1, only the up and down step 2 state ports were visible to the animal (the other ports were taped over), and the animal needed to learn that these ports deliver water. At stage 1.1, step 2 state ports were illuminated in a pseudorandom order with a 2-4s inter-trial interval without any auditory cues. Poking the illuminated port resulted in water reward delivery with 100% probability. When the animals completed >50 trials in one session at stage 1.1, they transitioned to stage 1.2 on the next session during which the auditory cues were introduced to signal active step 2 state ports and reward delivery. The delay in introducing auditory cues only at stage 1.2 was so that the animals do not get startled by the sounds such that they fail to explore the nose poke ports and get the water. When the animals completed >50 trials in one session in stage 1.2, they transitioned to stage 2 on the next session.

At the training stage 2, the step 1 choice ports were revealed to build the sequence of poking left/right and then up/down. All trials were forced-choice trials, so only one of the step 1 choice ports was illuminated on each trial, but the common/rare transition structure was used from the start. When the animals completed >50 trials in one session at stage 2, they transitioned to stage 3 on the next session.

At stage 3, the central port was revealed to complete the centre-left/right-up/down task sequence. Mice were switched to stage 4 when they completed >70 trials in one session.

At the fourth and final training stage, mice gradually learned how to find the “correct” choice based on different reward probabilities that change across blocks. Each daily session started in the same reward block that the previous session finished on. Free-choice trials and reward omissions were now introduced (i.e., not every choice was rewarded). Across the seven sub-stages of stage 4, the proportion of free-choice trials increased to 75%. Mice transitioned through sub-stages 4.1-4.6 if they completed >70 trials in one session. Mice transitioned to the final stage 4.7 when they were able to complete at least five reward reversals in a single session for at least three consecutive sessions. At stage 4.7, session length was increased to 90min and reward size reduced to 4µl to maximize the number of trials per session.

Animals were considered fully trained once they were consistently experiencing ≥6 reward reversals per session. Within each experiment, the animals were pseudo-randomly assigned to either the vehicle control or (±)-DOI group. Drug treatment was performed unblinded after the animals were fully trained, at the end of the last pre-treatment session, and behavioural testing resumed the next day.

When the same reversal problem is faced repeatedly, sophisticated automatization strategies may be developed which enable the animal to identify the relevant states of the world that have a fixed value. Serial reversals encourage automatized switching and planning for anticipated reward contingencies, and their underlying neural mechanism is different to situations when only one reversal is presented. This is why we anticipated that the novel reversal in transition probabilities would disrupt the subjects from using any habit-like strategies as the long-run predictive relationship between rewards and step 1 actions was broken.

*Exclusion criteria*

We initially trained 86 animals in total, and only four mice (one from Two-step Experiment 2 and three from Two-step experiment 3) had to be excluded because they did not reach the aforementioned criteria of performance (final $N$=82). These animals either failed to reach the final training stage 4.7 or consistently failed to do >5 reward reversals per session. One vehicle-treated animal in Two-step Experiment 1 had to be excluded from reward reversal adaptation analyses in the period after the transition reversal, as the subject never re-started doing >2 reward reversal per session from non-neutral reward blocks so not enough data was available to fit an average curve for that subject.

*Logistic regression analysis of choice behaviour*

All analysis was performed in Python (v.3.9.7) using both custom code and code adapted from Blanco-Pozo et. al. [3] and Akam et. al. [39].

Trial-to-trial learning and its modulation by drug treatment was assessed using a logistic regression model. The logistic regression was implemented using the scikit-learn function *linear_model.LogisticRegression* with the newton-cg solver. The dependent variable in logistic regression analyses was the animal's choices (excluding forced-choice trials) coded as stay (repeated choice) or switch (different choice). All sessions to be included in the regression analysis for one animal were concatenated such that each trial counted equally for each animal. The predictors were coded as a function of trial events as follows:

**Correct** (repeat correct choice): +0.5 for choosing an option which commonly leads to the step 2 state with higher reward probability (*correct choice*), -0.5 for choosing an option which commonly leads to the step 2 state with lower reward probability, 0 for any choices made during a neutral block. This predictor tracks the cumulative effect of past choices and outcomes to prevent correlations across trials from causing spurious loading on the *Transition* X *Outcome* interaction predictor [40].

**Choice** (repeat choice, side bias): +0.5 for repeated choices, -0.5 for different choices.

**Outcome** (repeat rewarded choice): +0.5 for rewarded trials, -0.5 for unrewarded trials.

**Transition** (repeat choices followed by a common transition): +0.5 for trials with a common transition, -0.5 for trials with a rare transition.

**Transition X Outcome** interaction (repeat common rewarded and rare unrewarded choices): +0.5 for rewarded trials with a common transition and unrewarded trials with a rare transition, -0.5 for unrewarded trials with a common transition and rewarded trials with a rare transition.

Notably, previous research in mice performing the two-step task showed that the influence of outcomes is asymmetrical – mice appear to be sensitive only to rewards and not to update their preferences following omissions [3]. The same asymmetry in learning rates from positive and negative feedback was observed in mice solving a simpler single-step probabilistic reinforcement learning task [41]. We show this here in the lagged regression analysis that signalled transitions affected future choices on rewarded but not unrewarded trials (Fig.2F).

Lagged logistic regression analysis assesses the subjects' choices as stay/shift as before, but now including the trail history at lags 1, 2, 3-4, 5-8, 9-12 (where the range of trials indicates the sum of the individual predictors over that range of lags) using the predictors:

**Common reward at lag *n***: +0.5 for repeated choice if the $n^{th}$ previous trial was rewarded via a common transition, 0 otherwise

**Rare reward at lag *n***: +0.5 for repeated choice if the $n^{th}$ previous trial was rewarded via a rare transition, 0 otherwise

**Common omission at lag *n***: +0.5 for repeated choice if the $n^{th}$ previous trial was not rewarded via a common transition, 0 otherwise

**Rare omission at lag *n***: +0.5 for repeated choice if the $n^{th}$ previous trial was not rewarded via a rare transition, 0 otherwise

When we wanted to determine if (±)-DOI and the added challenge of a novel transition reversal affected a mouse's drive to learn from omissions, we built a second logistic regression model that allowed us to evaluate the effect of transition type on the reinforcing effects of rewards and reward omissions separately. In addition to the *Correct*, *Choice*, and *Outcome* predictors as described above, this model included the following predictors:

**Reward by transition** (repeat rewarded choices with a common transition): +0.5 for rewarded trials with a common transition, -0.5 for rewarded trials with a rare transition, 0 for all unrewarded trials.

**Omission by transition** (repeat unrewarded choices with a common transition): +0.5 for unrewarded trials with a common transition, -0.5 for unrewarded trials with a rare transition, 0 for all rewarded trials.

*Simulations of Bayesian inference learning strategy*

In order to generate simulated data of subjects' expected choice strategies, we used Bayesian inference learning models described previously by Blanco-Pozo et al. [3]. The Bayesian inference strategy used a single variable to track the hidden state of the task (i.e., whether the up or down step 2 port has the higher reward probability). This variable was updated (i) using Bayes rule and (ii) based on the probability that a reward reversal has occurred (see Blanco-Pozo et al. [3] for the full details of the model). Note that in this model, both the rewards and the reward omissions in each step 2 state (up or down) were treated as different observations, and they contributed symmetrically to the state update (*Symmetric Bayesian Inference*). A variation of this model was the *Asymmetric Bayesian Inference* model which allowed for a differential update based on the type of outcome (rewards or omissions). Rewards in each step 2 state were treated as different observations (as in the Symmetric Bayesian Inference), but reward omissions in either step 2 state were treated as the same observation.

The chosen strategy was combined in a weighted sum with added bias and multi-trial perseveration parameters which modified the value of step 1 actions. A bias parameter increased the value of the left action by an amount determined by a bias strength parameter on all trials. A multi-trial perseveration parameter increased the value of the step 1 action chosen over the exponential moving average of earlier choices determined by an alpha multi-trial perseveration parameter. Each model was fit separately to subjects' actual behavioural data using maximum likelihood without priors. The optimisation was repeated 30 times starting with randomised initial parameter values drawn from a Beta distribution ($\alpha$=2, $\beta$=2) for unit range parameters, Gamma distribution ($\alpha$=2, $\beta$=0.4) for positive range parameters, and Normal distribution ($\sigma$=5) for unconstrained parameters. The best of these fits was used. To qualitatively compare data simulated from the model with real data, for each animal we simulated the same number of sessions (6 sessions) with their average number of trials per session (384.57 ±7.83 trials, mean ±SD), using parameter values from each animal's fits.

*Adaptation to reward reversals*

To compare the incorrect free-choices made in the first 20 trials after a reward reversal, firstly, per-subject means were calculated by averaging across all reversals from non-neutral reward blocks such that each reversal contributed equally. Then, the between-subject mean was calculated by averaging across individual subjects. Finally, we compared the between-subjects means of the two drug treatment groups using previously published methods [2, 39] to fit the between-subject mean choice probability trajectories with a double exponential decay function. The double exponential fit was calculated in Python, minimizing the error using the scikit-learn *minimize* function with the L-BFGS-B algorithm method. The starting value was determined by the mean choice probability in the final 10 trials before the reversal. The model was fitted to sessions from (±)-DOI- and vehicle-injected animals to give two sets of population level parameters:

$$\theta_{DOI} = \{\mu_{DOI}, \Sigma_{DOI}\} \text{ and } \theta_{VEH} = \{\mu_{VEH}, \Sigma_{VEH}\}$$

where $\theta_{DOI}$ are the parameters ($tau_{fast}$ and $tau_{slow}$) for trials from (±)-DOI - injected animals, and $\theta_{VEH}$ are the parameters ($tau_{fast}$ and $tau_{slow}$) for trials from vehicle-injected animals. The difference between the population means for the (±)-DOI and vehicle conditions was calculated as:

$$\Delta\mu_{true} = \mu_{DOI} - \mu_{VEH}$$

Permutation testing was used to assess the significance of differences in fits across (±)-DOI- and vehicle-treated animals (or between pre- and post-drug sessions of each treatment group separately). An ensemble of *N*=5000 permuted datasets was then created by shuffling the labels on sessions such that sessions were randomly assigned to the “(±)-DOI” and “vehicle” conditions (or “pre-drug” and “post-drug” conditions for within-subject comparisons across time). The double exponential was fit separately to sessions from (±)-DOI- and vehicle-injected animals for each permuted dataset, and the difference between population level means of $tau_{fast}$ and $tau_{slow}$ in the (±)-DOI and vehicle conditions was calculated for each permuted dataset *i* as:

$$\Delta\mu^{i}_{perm} = \mu^{i}_{DOI} - \mu^{i}_{VEH}$$

The distribution of $\Delta\mu_{perm}$ over the population of permuted datasets approximates the distribution under the null hypothesis that drug treatment does not affect the double exponential fit parameters ($tau_{fast}$ and $tau_{slow}$). The *P* values for the observed distances $\Delta\mu_{true}$ are then given by:

$$P = 2\min(\mathbf{M}/N, 1 - \mathbf{M}/N)$$

where **M** is the number of permutations (*N*) for which $\Delta\mu^{i}_{perm} > \Delta\mu_{true}$.

Note that the exponential time constants (*tau*) do not have a direct behavioural interpretation – they are simply the terms determining the shape of the exponential fit curve, i.e., how quickly the function decays. The lower the time constant, the faster the exponential decay. The two-phase model is therefore the sum of the fast and slow components ($tau_{fast/slow\ mix}$), each defined by their own rate constants, $tau_{fast}$ and $tau_{slow}$, respectively. Both phases are happening at all time points – it is not that the fast phase finishes and then the slow phase begins. Likewise, it is not that $tau_{fast}$ and $tau_{slow}$ reflect how long the system is in the fast or slow phase.

##### *Statistics*

*Nonlinear regression model fits and fit comparisons*

When testing for an effect of treatment on the change of a dependent variable *over time* we used nonlinear regression models instead of an analysis of variance (ANOVA) as ANOVAs test for a difference in means but do not consider any relationship between the data. ANOVA treats the different time points the same way it would treat different conditions. The fact that time points are sequential is ignored by ANOVA so the same results are obtained if the order of sessions is scrambled.

Nonlinear regression models were fitted in Prism v.9.4.0 (GraphPad Software Inc.) using unweighted least squares regression, considering each replicate $y$-value as an individual point, and evaluated using an extra sum-of-squares $F$ test. Outliers were identified with the ROUT method (Q=1%) and confidence intervals were calculated as asymmetrical profile likelihood. Normality and homoscedasticity of residuals was confirmed by the D'Agostino-Pearson omnibus and appropriate weighting test, respectively. Evidence of an inadequate model was evaluated by the replicates test for lack of fit. The main effect of time was signalled by the best-fit model not being a horizontal line ($H_0$: a straight-line fit across all post-reversal sessions). To test the drug effect, if a shared best-fit model was identified, an extra sum-of-squares $F$ test was used to determine if a shared global fit was sufficient ($H_0$), or if separate fits were warranted for each treatment group separately ($H_1$). We indicated the degree of evidence with the Akaike Information Criterion corrected for small sample sizes (AICc) to report the relative probabilities of the global fit or separate fits generating the data. If separate fits were suggested, we followed up these results by comparisons of individual versus shared model parameters. The drug effect was also indicated if the best-fits were from separate models for each treatment group (e.g., line and one-phase association), where the simpler model of the two served as $H_0$ (i.e., the line model was the null in the aforementioned example).

*General approach*

Significance of logistic regression coefficients was assessed using a one-sample *t*-test (or Wilcoxon signed-rank test for non-normally distributed data) comparing the distribution of the individual subjects' coefficients against zero. For lagged regression, Bonferroni multiple comparison correction was applied to a family of one predictor for all lags.

Significance of differences between pre-drug coefficients across treatment groups was assessed using an unpaired *t*-test, or a two-way repeated measures (RM) ANOVA in the case of a post-drug period. The number of trials, reward reversals, and correct choices pre-drug were compared using unpaired *t*-tests (or a two-way RM ANOVA in the case of a post-drug period).

For assessing the significant differences across sessions after transition reversal, nonlinear regression fits were used as described in the previous section.

All tests were two-sided with significance defined as $P<0.05$. Normal distribution of the data was verified with a D'Agostino-Pearson (omnibus K2) normality test and the normality of residuals was assessed in Q-Q plots. In the case of non-normal distributions, Wilcoxon signed-rank test and Mann-Whitney tests were used instead of the one-sample and unpaired *t*-tests, respectively. For repeated-measures, if Mauchly's test of sphericity was significant, Greenhouse-Geisser correction was applied. Data visualization and statistical analyses were done in Prism version 9.4.0 (GraphPad Software Inc.). Effect size calculations were done in JASP v.0.16.3 for MacOS Apple Silicon (JASP team). The experimenters were not blind to treatment group assignment during the analysis.

*Bayesian statistics*

We used Bayesian analysis to assess the strength of evidence to which the data supports the effects reported. Bayesian analyses were done in JASP v.0.16.3 for MacOS Apple Silicon. The Bayesian two-sided *t*-tests were implemented with the Cauchy prior (scale of 0.707). The implementation of Bayesian ANOVA was based on the *BayesFactor* package developed by Morey and Rouder [42]. The default uniform prior was used without enforcing the principle of marginality. To exclude random processes influencing the analyses and to ensure reproducibility of results, the seed was set to "123". Analysis of effects was reported in the form of inclusion Bayes factors ($BF_{incl}$) which give the odds ratio considering all models where the effect was included as $H_1$. All Bayes factors were reported as $BF_{10}$ to show evidence for the two-sided $H_1$ relative to $H_0$. When interpreting the evidence categories for Bayes factors, previously established guidelines [43] state the following: for the evidence in favour of $H_1$, the range of 1-3 is considered “weak”, the 3-10 range is considered “moderate”, and any values greater than 10 are considered “strong”; for the evidence in favour of $H_0$, the 0.33-1 range is considered “weak”, the 0.1-0.33 range is considered “moderate”, and values <0.1 are considered “strong”. The same evidence categories can be used to interpret the AICc test.

#### Supplementary Figures

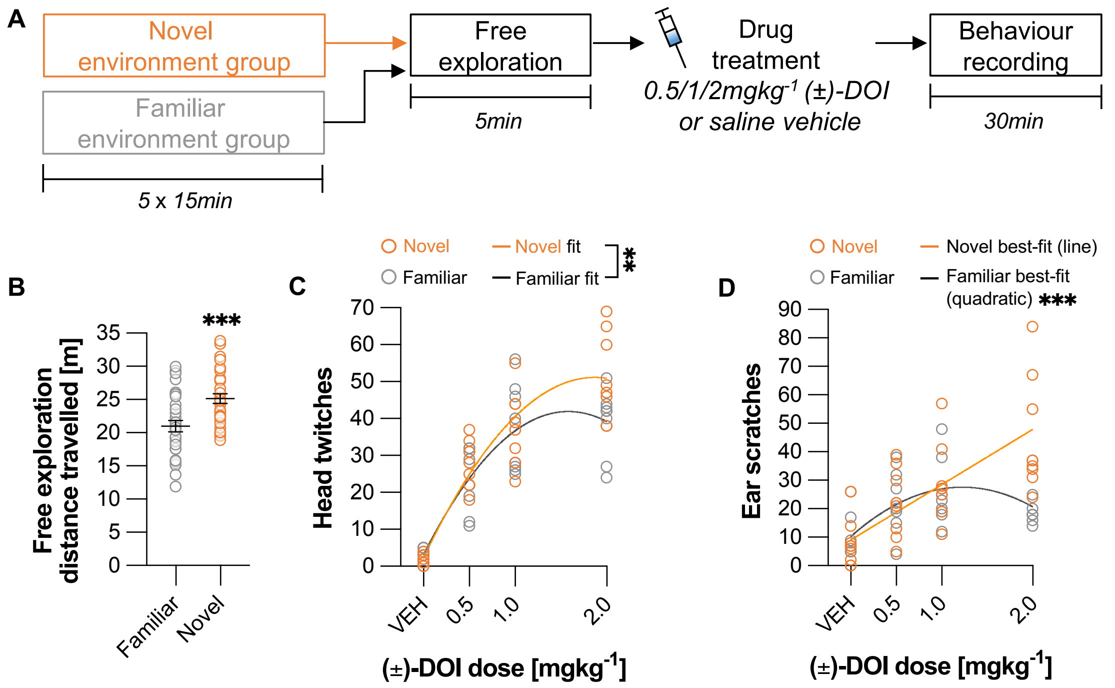


**Figure S1. Dose-dependent effects of (±)-DOI on the head-twitch and ear-scratch responses.**

**(A)** Experiment timeline. A subset of animals was habituated (5x15min) to the recording box before drug treatment (*familiar environment group*), while *the novel environment group* was exposed to the recording box only on the day of treatment. Both groups had 5min of free exploration before an injection of (±)-DOI (0.5, 1, or 2mgkg$^{-1}$) or saline vehicle. Animals were recorded for quantification of acute psychedelic-like responses in the first 30min after injection.

**(B)** In the 5min before the drug injection, habituated mice displayed significantly lower levels of locomotion compared to the mice being exposed to the recording box for the first time. Unpaired *t*-test, $t_{62}$= -3.71, *P*<0.001, Cohen's *d*= -0.93, $BF_{10}$=60.54. Data shown as mean ±SEM. $n_{group}$=32. ****P*<0.001.

**(C)** Dose-dependent increase in the number of head twitches followed the ascending limb of an inverted U-shaped curve. Nonlinear regression model of the total head twitches suggested separate fits for each environment. Comparison of quadratic Poisson fits *P*=0.002, likelihood ratio=15.2, AICc=59.5. The linear and quadratic terms were different across treatment environments (*P*=0.005), suggesting a later- and/or higher peaking model in the novel group. $n_{group}$=8. $N_{total}$=64. ***P*<0.01.

**(D)** Nonlinear Poisson fits of the total ear scratches required a different model for each environment. While the dose-response curve in the familiar environment preferred a quadratic fit ($P$<0.001, likelihood ratio=43.5, AICc>>100), with the highest 2mgkg$^{-1}$ dose being on the descending limb of the curve, a line fit was the best-fit for the novel environment (vs. quadratic fit $P$=0.054, likelihood ratio=3.71, AICc=1.89), suggesting the dose-response curve did not peak in the range of doses tested. $n_{group}$=8. $N_{total}$=64. ***$P$<0.001.

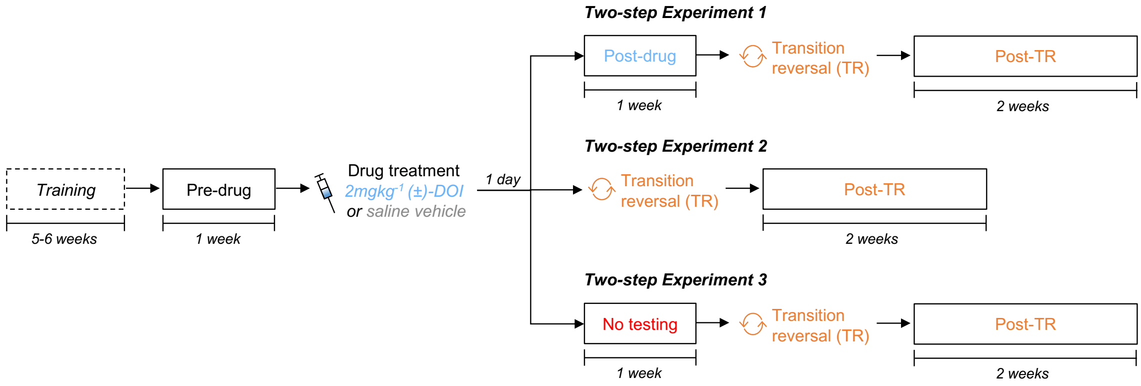


**Figure S2. Timelines used in two-step task experiments.** In all experiments, animals were treated with either saline vehicle or 2mgkg$^{-1}$ DOI when fully trained on the two-step task. In *Two-step Experiment 1*, the transition reversal (TR) was initiated after one week of testing the animals on an unchanged task in order to assess drug effects on cognitive adaptability with both the original task structure *and* with the reversed transition structure. In *Two-step Experiment 2*, the TR occurred immediately the next day after drug treatment in order to assess how time affects cognitive flexibility effects observed in the initial experiment. In *Two-step Experiment 3*, the TR was initiated one week after drug treatment but this time the animals were not tested on the original task during this first week in order to assess how the presence or absence of post-drug training affects cognitive flexibility effects observed in the initial experiment.

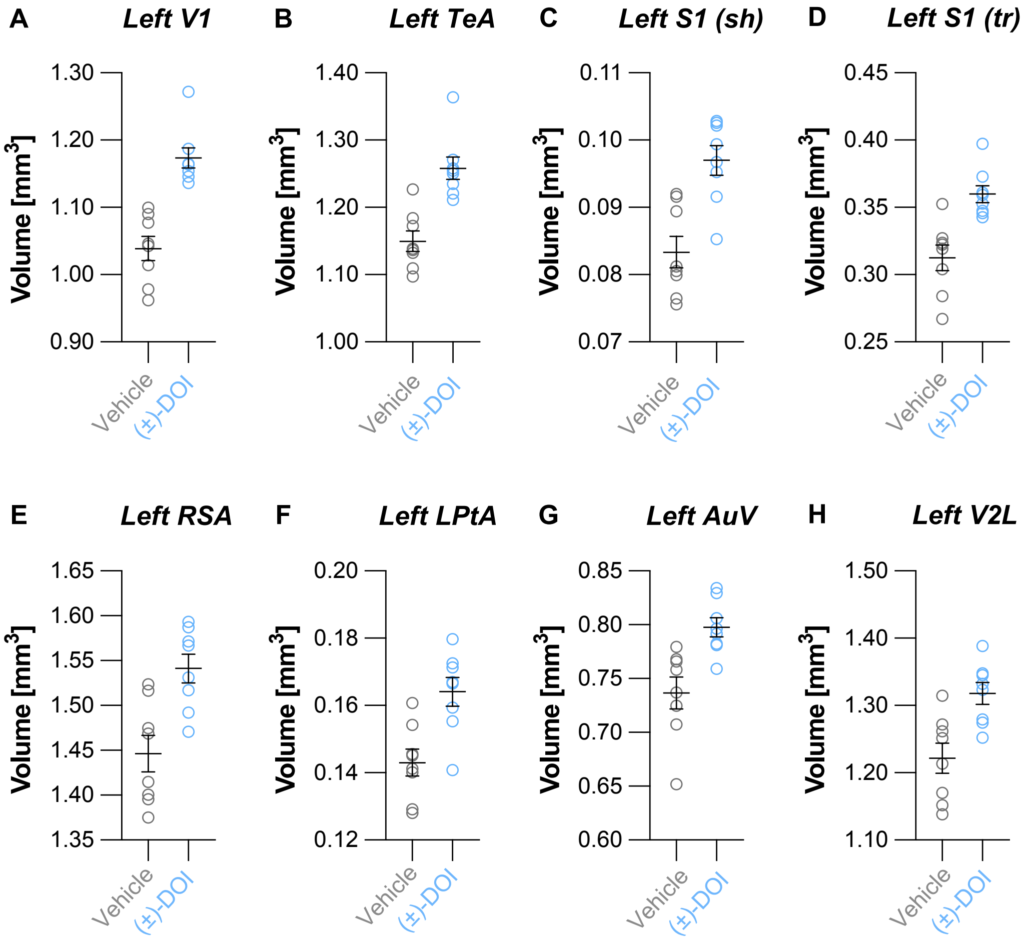


**Figure S3. A post-hoc visualization of ROIs that exhibited significant volume changes one day after (±)-DOI treatment.** ROIs shown were selected post-hoc based on a whole-brain analysis, so data are therefore shown for visualization purposes only, rather than for statistical inference. **(A)** Left primary visual area (V1). **(B)** Left temporal association area (TeA). **(C)** Left primary somatosensory area, shoulder region (S1(sh)). **(D)** Left primary somatosensory area, trunk region (S1(tr)). **(E)** Left retrosplenial agranular area (RSA). **(F)** Left lateral parietal association area (LPtA). **(G)** Left ventral secondary auditory area (AuV). **(H)** Left lateral secondary visual cortex (V2L). Data shown as mean ±SEM. $n_{group}$=8.

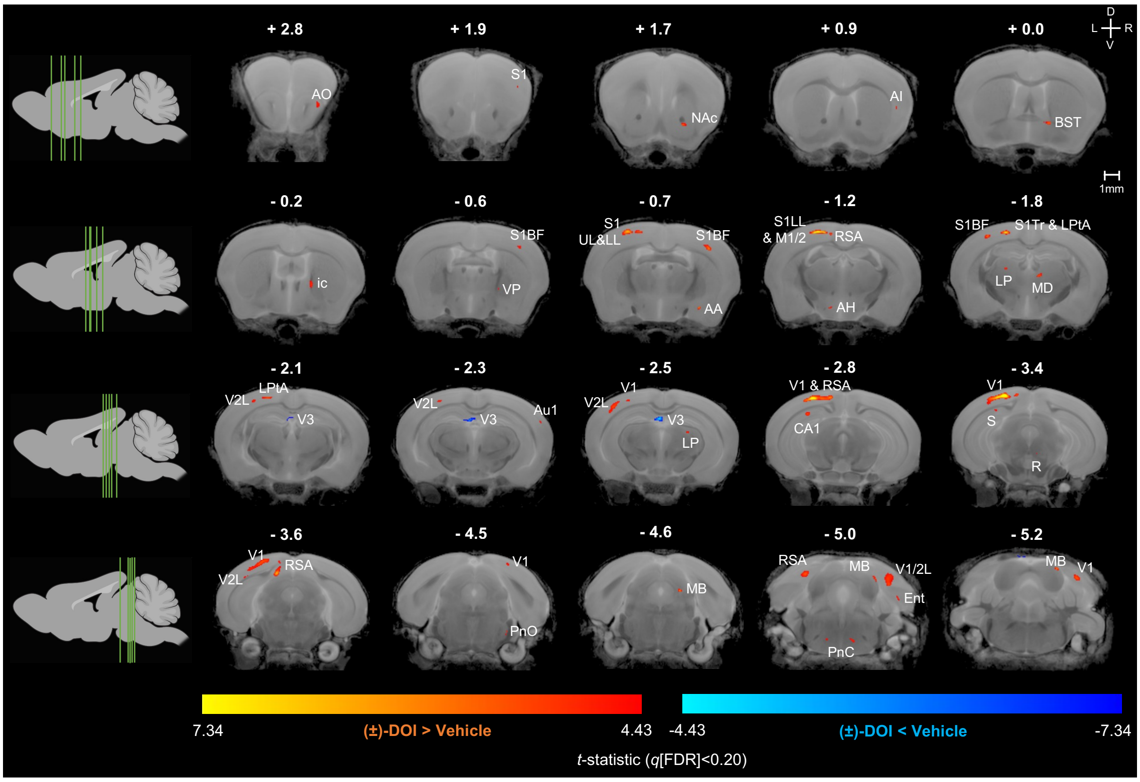


**Figure S4. Voxel-wise analysis of (±)-DOI-induced local changes in grey matter volume one day after treatment.** The False Discovery Rate (FDR) correction was applied to whole-brain voxel-wise linear model with an exploratory $q$<0.20 resulting in the marginal $t$-statistic threshold =4.43. All significant changes were volume increases ($t$>0), except for the third ventricle (V3). The maximum range of the colour legend represents the threshold $t$-statistic =7.34 at the strict $q$<0.05 to visualize which voxels pass the conservative criterion. $n_{group}$=8. D: dorsal. V: ventral. L: left. R: right. AA: anterior amygdaloid area. AH: anterior hypothalamic nuclei. AI: agranular insular area. AO: anterior olfactory nucleus. Au1: primary auditory cortex. BST: bed nucleus of stria terminalis. CA1: field CA1 of hippocampus. Ent: entorhinal cortex. ic: internal capsule. LP: lateral posterior thalamic nuclei. LPtA: lateral parietal association area. M1: primary motor area. M2: secondary motor area. MB: midbrain. MD: mediodorsal thalamic nuclei. NAc: nucleus accumbens. PnC: caudal pontine reticular nucleus. PnO: oral pontine reticular nucleus. R: red nucleus. RSA: retrosplenial area. S: subiculum. S1: primary somatosensory cortex; BF barrel field, UL upper limbs, LL lower limbs, Tr trunk. V1: primary visual cortex. V2L: lateral secondary visual cortex. VP: ventral pallidum.

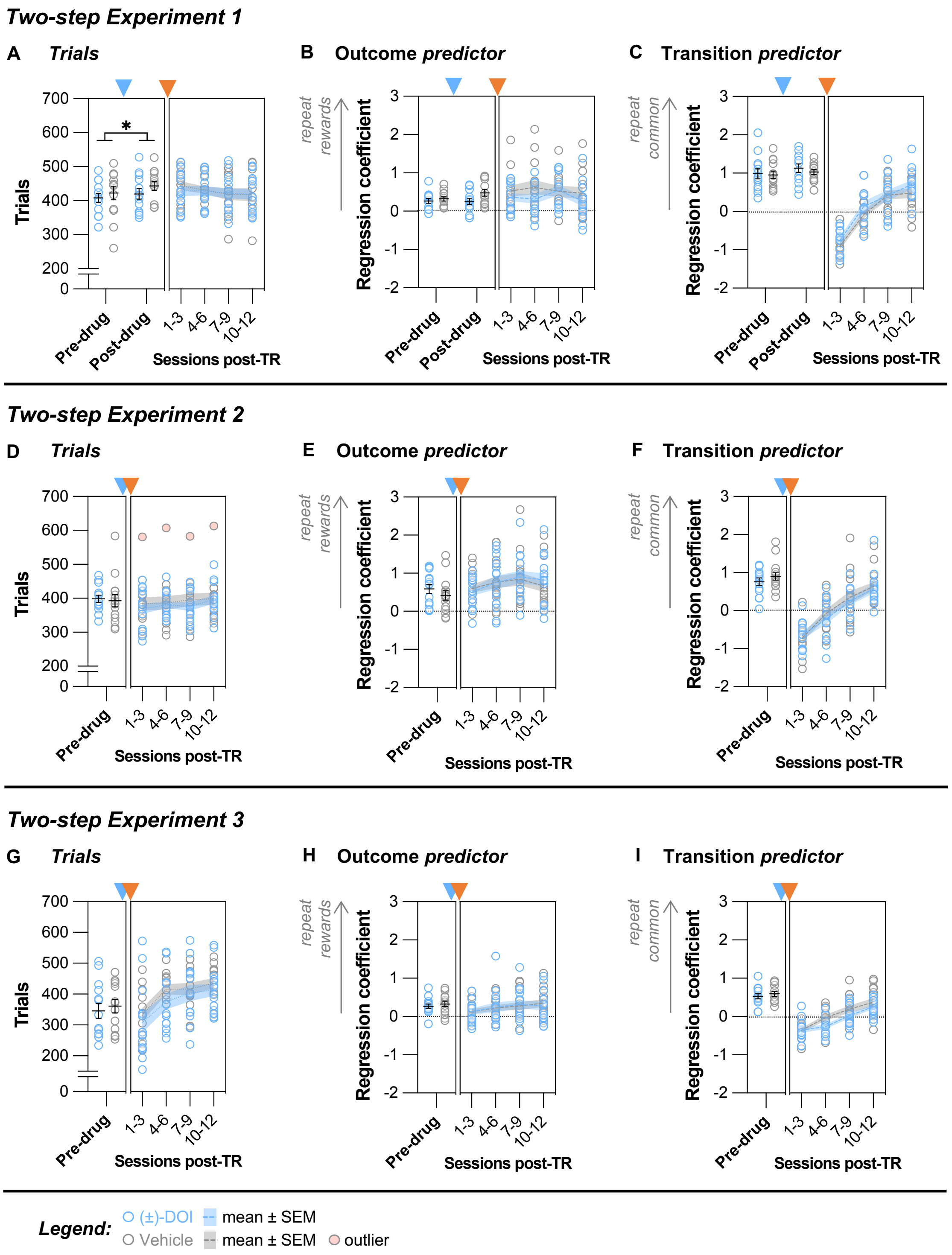


**Figure S5. Trial rates and simple reinforcing effects of rewards and common transitions across the experiments.**

5 **(A)** The number of trials the animals were completing per session did not differ across treatment groups before the drug treatment, but it increased marginally during the post-drug period irrespective of the type of treatment (two-way RM

ANOVA: Time $F_{1,24}$=6.4, $P$=0.018, $\eta_p^2$=0.21, $BF_{incl}$=3.03; Drug $F_{1,24}$=0.8, $P$=0.391, $BF_{incl}$=0.69; Time X Drug $F_{1,24}$=0.6, $P$=0.456, $BF_{incl}$=0.44). After the transition reversal (TR), the treatment groups remained comparable (nonlinear line regression fits $F_{2,100}$=0.24, $P$=0.786, $AICc_{10}$=0.14, $slope_{global}$=-6.4, CI [-15.8, 2.9]).

**(B)** The logistic regression model predictor for *Outcome* was not significantly different before or after injection across animals pre-selected for vehicle or (±)-DOI treatment (Pre- vs. Post-drug two-way RM ANOVA: Time $F_{1,24}$=2.1, $P$=0.161, $BF_{incl}$=0.62; Drug $F_{1,24}$=2.5, $P$=0.127, $BF_{incl}$=0.97; Time X Drug $F_{1,24}$=3.3, $P$=0.082, $BF_{incl}$=1.12). TR did not affect how trial outcome influenced subsequent choices in either treatment condition (nonlinear line regression fits $F_{2,100}$=1.28, $P$=0.283, $AICc_{10}$=0.42, $slope_{global}$= -0.02, CI [-0.11, 0.07]).

**(C)** The logistic regression model predictor for *Transition* was not significantly different before or after injection across animals pre-selected for vehicle or (±)-DOI treatment (Pre- vs. Post-drug two-way RM ANOVA: Time $F_{1,24}$=3.8, $P$=0.063, $BF_{incl}$=1.23; Drug $F_{1,24}$=0.2, $P$=0.624, $BF_{incl}$=0.53; Time X Drug $F_{1,24}$=1.5, $P$=0.223, $BF_{incl}$=0.41). The *Transition* predictor switched signs after TR, indicating that the animals were initially still following the pre-reversal transition structure, repeating choices followed by previously common but now rare transitions. They learned how to reverse their strategy over time, as indicated by the positive *Transition* loading by post-TR sessions 7-9, but this reversal was not different across treatment groups (nonlinear one-phase association regression fits $F_{3,98}$=1.06, $P$=0.372, $AICc_{10}$=0.18, $tau_{global}$=1.27, CI [0.76, 2.52]).

**(D)** The number of trials the animals were completing per session did not differ across treatment groups before the drug treatment (unpaired *t*-test $t_{27}$=-0.26, $P$=0.798, $BF_{10}$=0.36) or after TR (nonlinear line regression fits $F_{2,108}$=0.18, $P$=0.840, $AICc_{10}$=0.14, $slope_{global}$=6.6, CI [-1.1, 14.4]).

**(E)** The logistic regression model predictor for *Outcome* was not significantly different across animals pre-selected for vehicle or (±)-DOI treatment (unpaired *t*-test $t_{27}$=-1.03, $P$=0.313, $BF_{10}$=0.52). TR did not affect how trial outcome influenced subsequent choices in either treatment condition (nonlinear line regression fits $F_{2,111}$=0.75, $P$=0.474, $AICc_{10}$=0.23, $slope_{global}$=0.06, CI [-0.03, 0.15]).

**(F)** The logistic regression model predictor for *Transition* was not significantly different across animals pre-selected for vehicle or (±)-DOI treatment (unpaired *t*-test $t_{27}$=1.00, $P$=0.327, $BF_{10}$=0.51). Strategy reversal after TR was not different across treatment groups (nonlinear line regression fits $F_{2,112}$=0.32, $P$=0.726, $AICc_{10}$=0.16, $slope_{global}$=0.43, CI [0.36, 0.51]).

**(G)** The number of trials the animals were completing per session did not differ across treatment groups before the drug treatment (unpaired *t*-test $t_{25}$=0.475, $P$=0.639, $BF_{10}$=0.39) or after TR (nonlinear line regression fits $F_{2,104}$=1.20, $P$=0.306, $AICc_{10}$=0.39). The trial rate was initially lower after the week of no testing and increased over time after the TR ($slope_{global}$=33.1, CI [18.4, 47.8]).

**(H)** The logistic regression model predictor for *Outcome* was not significantly different across animals pre-selected for vehicle or (±)-DOI treatment (Pre-drug period, unpaired *t*-test $t_{25}$=0.63, $P$=0.536, $BF_{10}$=0.42). TR did not affect how trial outcome influenced subsequent choices in either treatment condition (nonlinear line regression fits $F_{2,104}$=0.065, $P$=0.937, $AICc_{10}$=0.12, $slope_{global}$=0.06, CI [-0.002, 0.12]).

**(I)** The logistic regression model predictor for *Transition* was not significantly different across animals pre-selected for vehicle or (±)-DOI treatment (Pre-drug period, unpaired *t*-test $t_{25}$=0.69, $P$=0.497, $BF_{10}$=0.43). Strategy reversal after TR was not different across treatment groups (nonlinear line regression fits $F_{2,104}$=2.98, $P$=0.055, $AICc_{10}$=2.28, $slope_{global}$=0.23, CI [0.18, 0.28]).

Data shown as mean ±SEM. Experiment 1: $n_{Veh}$=13. $n_{(\pm)\text{-DOI}}$=13. Experiment 2: $n_{Veh}$=15. $n_{(\pm)\text{-DOI}}$=14. Experiment 3: $n_{Veh}$=13. $n_{(\pm)\text{-DOI}}$=14.

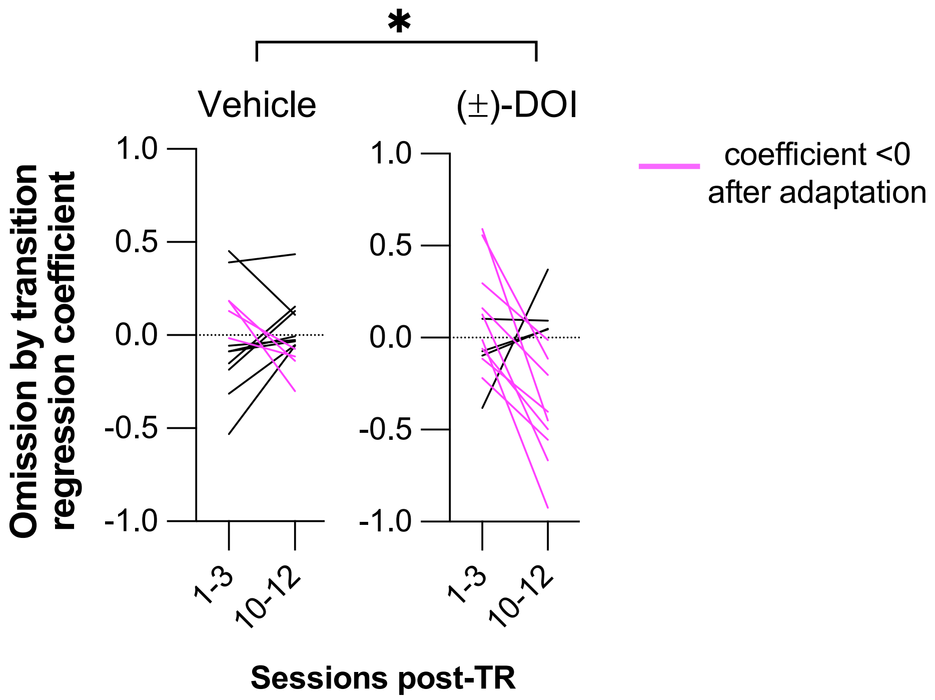


**Figure S6. More of the (±)-DOI-treated than vehicle-treated Experiment 1 mice had a lower *Omission by transition* regression coefficient by the end of the transition reversal adaptation period.** Tracking the regression coefficient from the beginning to end of the adaptation to the transition reversal (TR), the magenta line indicates the animals whose coefficient dropped below zero (or became more negative, for those animals which already had a negative coefficient at the start). Only 4 out of 13 mice treated with saline vehicle had coefficients drop to values <0, compared to 9 out of 13 mice treated with (±)-DOI (likelihood ratio test =3.95, *P*=0.047). The two treatment groups had comparable numbers of animals that had a negative coefficient at the start of the adaptation period (8 of 13 in vehicle group, 7 of 13 in (±)-DOI group, likelihood ratio test =0.16, *P*=0.691, $BF_{10}$=2.80).

Data shown as mean ±SEM. $n_{Veh}$=13. $n_{(\pm)\text{-}DOI}$=13.

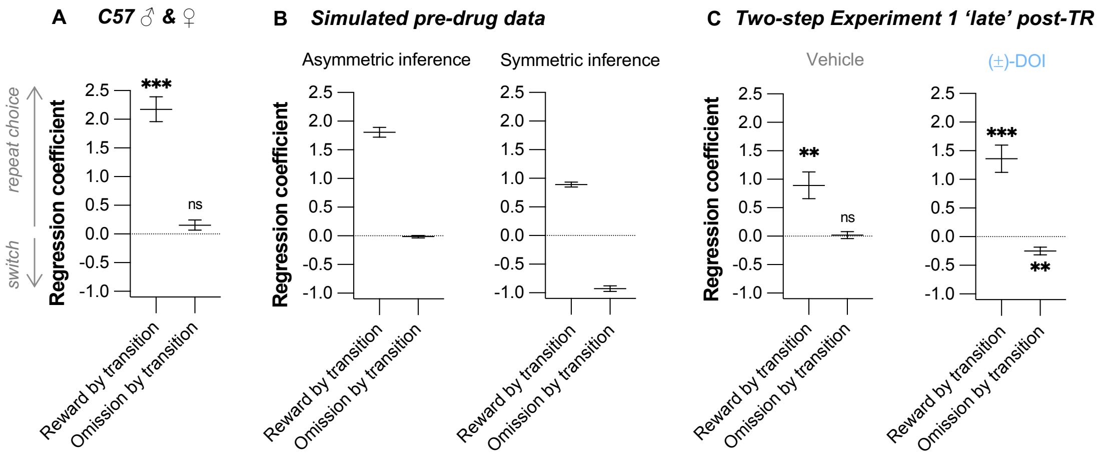


**Figure S7. In general, trial-by-trial choice strategy of C57 mice is consistent with asymmetric inference learning, but (±)-DOI-treated Experiment 1 animals shifted towards a more symmetric inference learning.**

**(A)** We reanalysed data from a previous report [3] on wild-type and DAT-Cre C57 male and female mice solving the two-step task without any experimental manipulation using our logistic regression model that looks at the effect of transition type on rewards and reward omissions separately. We confirmed the choice strategy was comparable to the one we observed in our experiments. The loading on the *Omission by transition* predictor is not significantly different from 0, i.e., these animals also did not use reward omission experience as informative of task state. One sample *t*-tests against zero: *Reward by transition* $t_{17}$=10.03, $P$<0.001, Cohen's $d$ =2.36, $BF_{10}$>>100; *Omission by transition* $t_{17}$=1.77, $P$=0.095, $BF_{10}$=0.88. $n$=18.

**(B)** Asymmetric inference has been reported as the best-fit model explaining mouse behaviour in the two-step task whereby mice use inference to update their belief of the task state but using information from rewards only, and not from reward omissions [3]. We simulated data for both asymmetric and symmetric inference learning based on the pre-drug performance of mice from all three of our experiments. The loading on the *Omission by transition* predictor was expected to be minimal in the asymmetric inference model. With the symmetric inference model, rewards and reward omissions would be treated as equally informative. The resulting pattern reflects a tendency to repeat rewarded choices preceded by common transitions while switching away from omission trials with a common transition. Our data shown in this report and the previously published data shown in the previous panel are both qualitatively more comparable to the asymmetric inference learning model, as predicted. $n$=82.

**(C)** In the Experiment 1 mice, which underwent a novel transition reversal one week after drug injection, the *Omission by transition* predictor shifted to being significantly negative in the second week of adaptation ('late' post-TR period) only for the (±)-DOI-treated mice. One sample *t*-tests against zero: Vehicle *Reward by transition* $t_{12}$=3.80, *P*=0.003, Cohen's *d*=1.05, $BF_{10}$=18.1, *Omission by transition* $t_{12}$=0.29, *P*=0.773, $BF_{10}$=0.29; (±)-DOI *Reward by transition* $t_{12}$=5.74, *P*<0.001, Cohen's *d*=1.59, $BF_{10}$=307.3, *Omission by transition* $t_{12}$= -3.76, *P*=0.003, Cohen's *d* = -1.04, $BF_{10}$=17.0. We note that the behavioural pattern of Experiment 1 (±)-DOI-treated mice after the transition reversal, specifically the negative loading on the *Omission by transition* predictor, is closer to the symmetrical inference learning model than the behavioural pattern of vehicle-treated mice. $n_{group}$=13.

Data shown as mean ±SEM. ***$P$<0.001. **$P$<0.01.

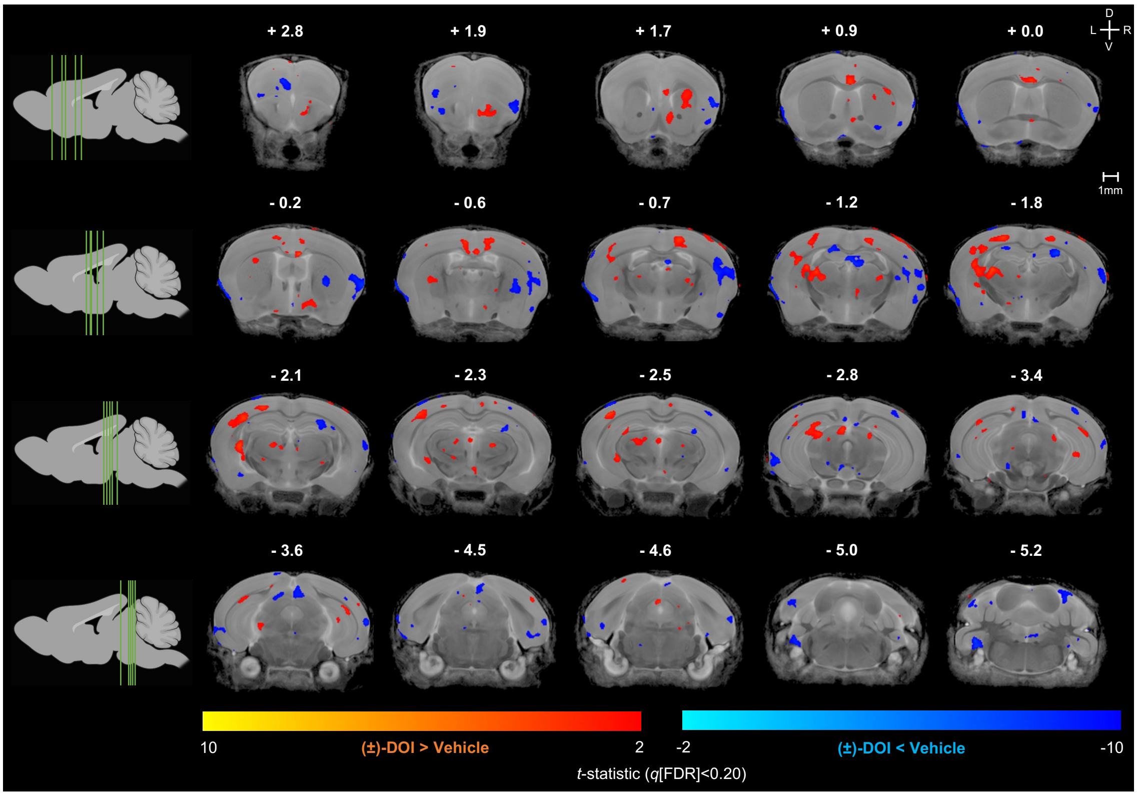


**Figure S8. Voxel-wise analysis of (±)-DOI-induced local changes in grey matter volume three weeks after treatment.** The False Discovery Rate (FDR) correction was applied to whole-brain voxel-wise linear model with an exploratory $q<0.20$ resulting in no significant findings for the main effect of drug treatment. Unthresholded map visualizes voxel changes near an approximate marginal *t*-statistic threshold. Data are therefore shown for visualization purposes only, rather than for statistical inference. $n_{group}=10$. D: dorsal. V: ventral. L: left. R: right.

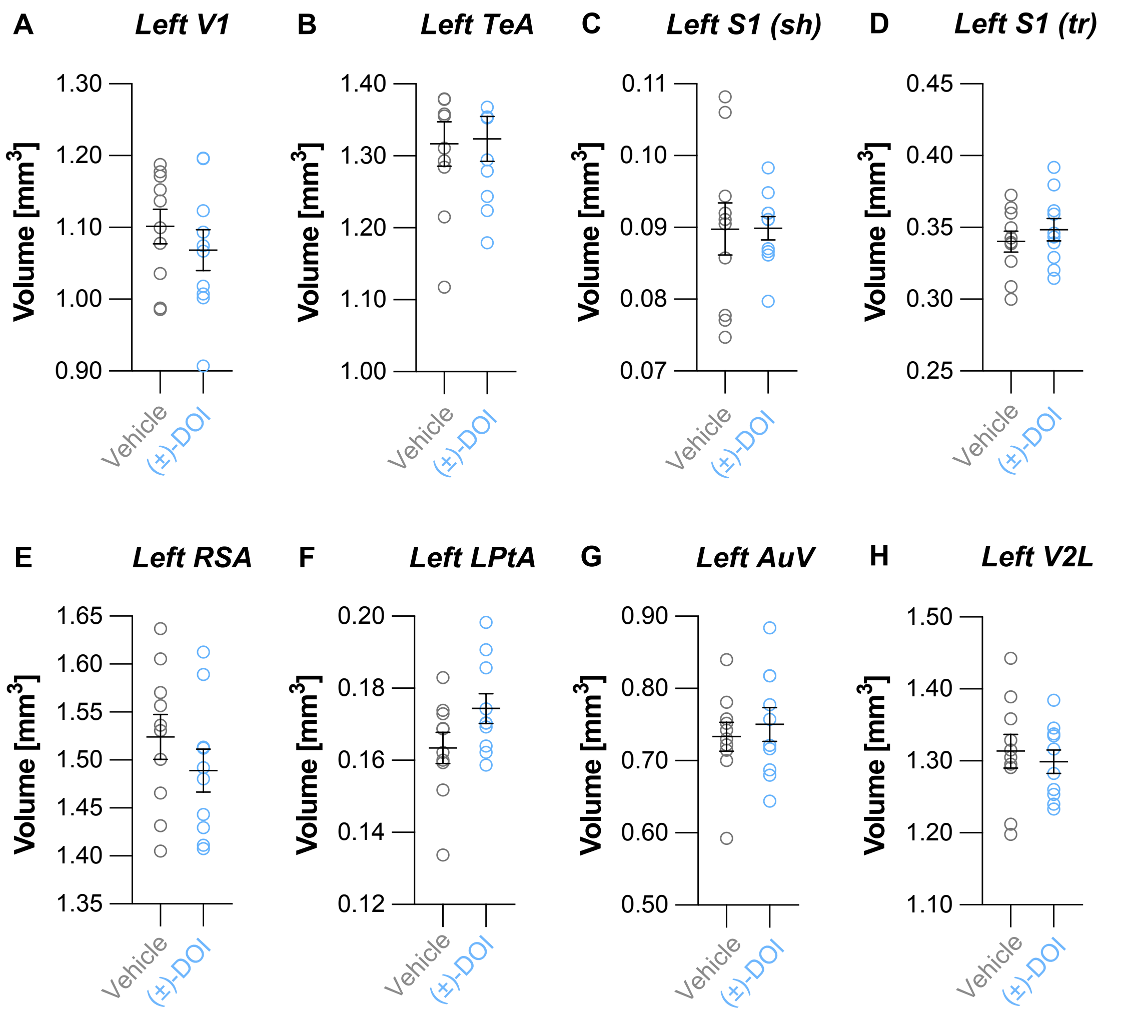


**Figure S9. In brains taken three weeks after treatment, ROIs that had exhibited significant volume changes in the initial study, in brains sampled one day after (±)-DOI treatment, were comparable across treatment groups.** ROIs shown were selected post-hoc based on a whole-brain analysis, so data are therefore shown for visualization purposes only, rather than for statistical inference. **(A)** Left primary visual area (V1). **(B)** Left temporal association area (TeA). **(C)** Left primary somatosensory area, shoulder region (S1(sh)). **(D)** Left primary somatosensory area, trunk region (S1(tr)). **(E)** Left retrosplenial agranular area (RSA). **(F)** Left lateral parietal association area (LPtA). **(G)** Left ventral secondary auditory area (AuV). **(H)** Left lateral secondary visual cortex (V2L). Data shown as mean ±SEM. $n_{group}$=10.

#### Supplementary Tables

**Table S1. The training stages for the two-step task.** Grey-shaded areas indicate when task conditions changed. Operant box nose poke ports: up (U), down (D), left (L), right (R), centre (C).

| Stage | Learning outcome | Available Pokes | Transition Probability *-Common-* | Transition Probability *-Rare-* | Reward Probability *-Common-* | Reward Probability *-Rare-* | Reward Probability *-Neutral-* | % Free Choice | Reward Volume [µL] | Learning criterion | Sessions to criterion (min – max) | Change from previous stage |
|---|---|---|---|---|---|---|---|---|---|---|---|---|
| **1.1** | U/D pokes deliver reward | **UD** | **NA** | **NA** | **NA** | **NA** | **1** | **0%** | 15 | >50 trials | 1 | |
| **1.2** | | **UD** | **NA** | **NA** | **NA** | **NA** | **1** | **0%** | 12 | >50 trials | 1 - 2 | auditory cue (different frequency for up and down) before reward is delivered |
| **2** | L/R → U/D sequence | **UDLR** | **0.8** | **0.2** | **1** | **1** | **1** | **0%** | 12/10 | >50 trials | 1 - 5 | left & right ports, auditory cues with different frequency for transitions with 80/20% common/rare probabilities |
| **3** | C → L/R → U/D sequence | **UDLRC** | **0.8** | **0.2** | **1** | **1** | **1** | **0%** | 12/10 | >70 trials | 1 - 3 | initiation (centre) port |
| **4.1** | finding the 'correct' choice based on different reward probabilities that reverse across blocks | **UDLRC** | **0.8** | **0.2** | **0.9** | **0.7** | **0.8** | **0%** | 12/10 | >70 trials | 1 | reward probabilities, block switch after 20-30 trials, white noise signals no reward |
| **4.2** | | **UDLRC** | **0.8** | **0.2** | **0.9** | **0.5** | **0.7** | **25%** | 10 | >70 trials | 1 - 2 | 25% free choice trials, rare/neutral reward probabilities |
| **4.3** | | **UDLRC** | **0.8** | **0.2** | **0.9** | **0.3** | **0.6** | **25%** | 10 | >70 trials | 1 | rare/neutral reward probabilities |
| **4.4** | | **UDLRC** | **0.8** | **0.2** | **0.9** | **0.1** | **0.5** | **25%** | 10 | >70 trials | 1 | rare/neutral reward probabilities |
| **4.5** | | **UDLRC** | **0.8** | **0.2** | **0.9** | **0.1** | **0.5** | **50%** | 10 | >70 trials | 1 | 50% free choice trials, block switch in non-neutral blocks triggered by animal's behaviour |
| **4.6** | | **UDLRC** | **0.8** | **0.2** | **0.9** | **0.1** | **0.5** | **75%** | 10/8/6 | >5 blocks for 3 consecutive sessions | 4 - 16 | 75% free choice trials, 90min session length for slow learners |
| **4.7** | | **UDLRC** | **0.8** | **0.2** | **0.8** | **0.2** | **0.5** | **75%** | 6/4 | ≥6 blocks in >6 sessions | 1 - 20 | common/rare reward probabilities, 90min session length |

**Table S2. Regional volume differences one day after 2mg/kg (±)-DOI treatment.** The false discovery rate (FDR) threshold was set to $q<0.20$. The first column are the regions of interest (ROI) with statistically significant differences in volume between the (±)-DOI-treated and saline vehicle-treated samples. The following columns are the *F*-statistics of the significance of the structure, the drug term marginal *t*-statistics and corresponding FDR-corrected *q*-value. The final columns state the mean volumes of each ROI with the 95% confidence intervals (CIs) shown as [lower limit, upper limit], as well as the approximate volume change in (±)-DOI-treated samples compared to the controls. The degrees of freedom are given in brackets. V1: primary visual area. TeA: temporal association area. S1: primary somatosensory area. RSA: retrosplenial agranular area. LPtA: lateral parietal association area. AuV: ventral secondary auditory area. V2L: lateral secondary visual area.

| | | | | *VEH* | | *DOI* | | |
|---|---|---|---|---|---|---|---|---|
| **ROI** | $F_{(1,14)}$ | $t_{(14)}$ | $q$ **value** | **Mean [mm$^3$]** | **95% CI** | **Mean [mm$^3$]** | **95% CI** | ***% change*** **± SEM** |
| *Left V1* | 33.06 | 5.75 | 0.016 | 1.039 | [0.996, 1.081] | 1.174 | [1.138, 1.209] | 13.0 ± 2.3 |
| *Left TeA* | 23.33 | 4.83 | 0.043 | 1.150 | [1.114, 1.185] | 1.258 | [1.219, 1.297] | 9.4 ± 2.0 |
| *Left S1 (shoulder)* | 18.02 | 4.25 | 0.079 | 0.083 | [0.078, 0.089] | 0.097 | [0.092, 0.102] | 16.4 ± 3.9 |
| *Left S1 (trunk)* | 17.23 | 4.15 | 0.079 | 0.313 | [0.290, 0.335] | 0.360 | [0.345, 0.374] | 15.2 ± 3.6 |
| *Left RSA* | 13.53 | 3.68 | 0.145 | 1.446 | [1.398, 1.494] | 1.541 | [1.503, 1.579] | 6.6 ± 1.8 |
| *Left LPtA* | 13.05 | 3.61 | 0.145 | 0.143 | [0.134, 0.152] | 0.164 | [0.154, 0.174] | 14.7 ± 4.1 |
| *Left AuV* | 12.47 | 3.53 | 0.145 | 0.737 | [0.700, 0.772] | 0.798 | [0.777, 0.819] | 8.3 ± 2.3 |
| *Left V2L* | 12.19 | 3.49 | 0.145 | 1.222 | [1.168, 1.274] | 1.318 | [1.280, 1.356] | 7.9 ± 2.3 |

14. Odland AU, Jessen L, Kristensen JL, Fitzpatrick CM, Andreasen JT. The 5-hydroxytryptamine 2A receptor agonists DOI and 25CN-NBOH

decrease marble burying and reverse 8-OH-DPAT-induced deficit in spontaneous alternation. Neuropharmacology. 2019:107838.

15. Goldberger L. Cognitive test performance under LSD-25, placebo and isolation. Journal of Nervous and Mental Disease. 1966. 1966.

21. dos Santos RG, Osório FL, Crippa JAS, Riba J, Zuardi AW, Hallak JEC. Antidepressive, anxiolytic, and antiaddictive effects of ayahuasca, psilocybin and lysergic acid diethylamide (LSD): a systematic review of

clinical trials published in the last 25 years. Ther Adv Psychopharmacol. 2016;6:193–213.

28. Lerch JP, Yiu AP, Martinez-Canabal A, Pekar T, Bohbot VD, Frankland PW, et al. Maze training in mice induces MRI-detectable brain shape

changes specific to the type of learning. Neuroimage. 2011;54:2086–2095.

36. Daw ND, Dayan P. The algorithmic anatomy of model-based evaluation. Philosophical Transactions of the Royal Society B: Biological Sciences. 2014;369.

37. Doll BB, Duncan KD, Simon DA, Shohamy D, Daw ND. Model-based choices involve prospective neural activity. Nat Neurosci. 2015;18:767.

38. Akam T, Lustig A, Rowland JM, Kapanaiah SKT, Esteve-Agraz J, Panniello M, et al. Open-source, Python-based, hardware and software for controlling behavioural neuroscience experiments. Elife. 2022;11.

39. Akam T, Rodrigues-Vaz I, Marcelo I, Zhang X, Pereira M, Oliveira RF, et al. The Anterior Cingulate Cortex Predicts Future States to Mediate Model-Based Action Selection. Neuron. 2021;109:149-163.e7.

40. Akam T, Costa R, Dayan P. Simple plans or sophisticated habits? State, transition and learning interactions in the two-step task. PLoS Comput Biol. 2015;11:e1004648.

41. Cieślak PE, Ahn W-Y, Bogacz R, Parkitna JR. Selective effects of the loss of NMDA or mGluR5 receptors in the reward system on adaptive decision-making. ENeuro. 2018;5.

42. Morey RD, Rouder JN. Package ‘bayesfactor’. 2015. https://richarddmorey.github.io/BayesFactor/. Accessed 2 December 2022.

43. van Doorn J, van den Bergh D, Böhm U, Dablander F, Derks K, Draws T, et al. The JASP guidelines for conducting and reporting a Bayesian analysis. Psychon Bull Rev. 2021;28:813–826.
